# KINNTREX: A Neural Network Unveils Protein Mechanism from Time Resolved X-ray Crystallography

**DOI:** 10.1101/2023.10.06.561268

**Authors:** Gabriel Biener, Tek Narsingh Malla, Peter Schwander, Marius Schmidt

## Abstract

Here, a machine learning method based on a kinetically informed neural network (NN) is introduced. The proposed method is designed to analyze a time series of difference electron density (DED) maps from a time-resolved X-ray crystallographic experiment. The method is named KINNTREX (Kinetics Inspired NN for Time-Resolved X-ray Crystallography). To validate KINNTREX, multiple realistic scenarios were simulated with increasing level of complexity. For the simulations, time-resolved X-ray data was generated that mimic data collected from the photocycle of the photoactive yellow protein (PYP).

KINNTREX only requires the number of intermediates and approximate relaxation times (both obtained from a singular valued decomposition) and does not require an assumption of a candidate mechanism. It successfully predicts a consistent chemical kinetic mechanism, together with difference electron density maps of the intermediates that appear during the reaction. These features make KINNTREX attractive for tackling a wide range of biomolecular questions. In addition, the versatility of KINNTREX can inspire more NN-based applications to time-resolved data from biological macromolecules obtained by other methods.

## 1. Introduction

Biological macromolecules perform essential functions in living organisms. One class of these molecules, proteins, is of particular importance. Proteins are involved in all functions of life, spanning from light perception to the catalysis of essential reactions. To perform their function, proteins must undergo structural (conformational) changes. For example, a catalytically active protein, an enzyme, changes its conformation upon binding of a substrate, during the catalytic conversion of the substrate to product, and when the product leaves (Cornish-Bowden, 2004). Distinct conformational states along the reaction pathway are called intermediates. The sequence of transitions between intermediates is described by a scheme referred to as a chemical, kinetic mechanism (Steinfeld *et al*., 1999). Fig. 1 shows several examples of these mechanisms. Mechanisms may include irreversible processes indicated by single arrows (see Fig. 1) or reversible processes (arrows pointing in both directions) where an equilibrium between two intermediates is established.

**Figure 1.**
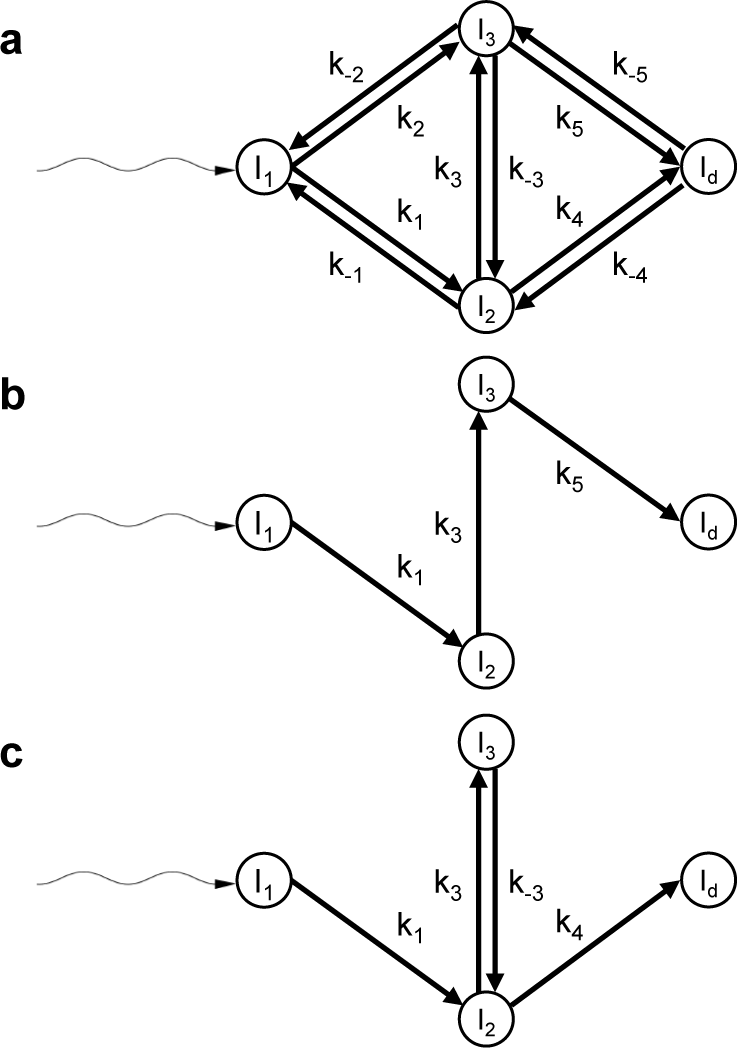
Chemical kinetic mechanisms with three intermediates for the (a) general case, (b) an irreversible sequential mechanism, and (c) a dead-end mechanism for a bio-macromolecular reaction initiated by light (wavy arrows). Circles represent intermediate states. The straight arrows labeled with reaction rate coefficients (RRCs) denote transitions between intermediates.

Multiple methods have been developed to determine protein structures with near atomic resolution. X-ray crystallography (Blake *et al*., 1965), cryogenic electron microscopy (Yip *et al*., 2020), and nuclear magnetic resonance (Wüthrich, 1990) all offer static snapshots of the protein structure. As a protein changes its structure along the reaction pathway, a single structure is not sufficient to comprehensively describe its function. Time resolved X-ray crystallography (TRX) aims at determining structure and dynamics at the same time (Moffat, 2001). TRX captures X-ray diffraction patterns during the time the protein performs its function. These diffraction patterns are then processed to yield time-dependent electron density maps (Schmidt, 2019).

Typically, only a small fraction of molecules in a crystal participates in a reaction because methods to initiate a reaction can be quite ineffective (Srajer & Schmidt, 2017). As a result, the extent of reaction initiation can be small and on the order of < 10 %. Conventional electron density maps are insensitive to the presence of a small admixture of reacting molecules in the presence of a large amount of protein at rest. However, with difference electron density (DED) maps even small amounts of reacting molecules can be detected. A DED map is obtained by subtracting a reference electron density, where all molecules are at rest, from the time-dependent electron density (Henderson & Moffat, 1971). The DED map has positive and negative features. The negative features are found at locations in the reference structure where an atom has moved away. Positive features are found at positions where the atoms have moved to, or when additional atoms, e.g., those of a ligand, bind. The biomolecular reaction is then probed by a time-series of difference maps calculated from TRX data best collected at equidistant time-points along logarithmic time (Moffat, 2001, Schmidt *et al*., 2003, Rajagopal *et al*., 2004, Ihee *et al*., 2005, Schmidt *et al*., 2013, Schotte *et al*., 2012).

The DED map at each measured time point is a sum of all intermediate states, each multiplied by a concentration value at that time point (Schmidt *et al*., 2003, Moffat, 2001). The time-dependent concentrations of the intermediates are determined by the chemical, kinetic mechanism and the rate coefficients that characterize the transitions between the intermediates (Moffat, 1989, 2001). The time evolution of the concentrations of an intermediate during a reaction is called a concentration profile. Fig. 2 shows such concentration profiles calculated from two mechanisms, the sequential (Fig. 1b) and dead-end (Fig. 1c) mechanisms. Two sets of reaction rate coefficients (RRC) are used for each case. It is evident that at almost all time points the signal is generated by a mixture of species at different concentrations. The concentration profiles are calculated by solving the coupled differential equations of the kinetic mechanism (Steinfeld *et al*., 1999).

**Figure 2.**
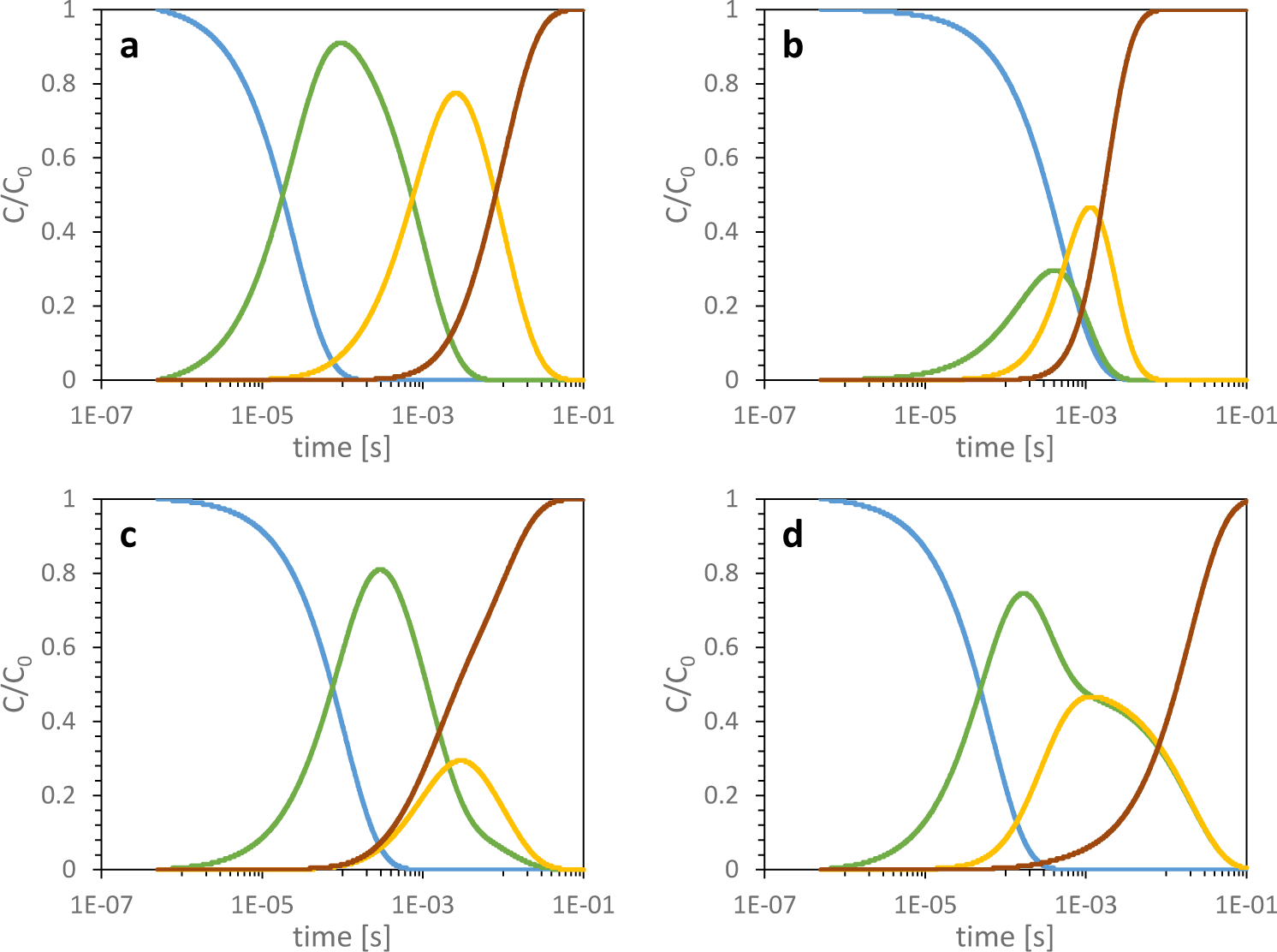
Concentration profiles derived from simulations that mimic the reaction of the blue light photoreceptor photoactive yellow protein. Fractional concentrations C/C_0_ are plotted as a function of log t. Concentrations of intermediates *I*_*1*_ to *I*_*3*_ are represented by blue, green, and yellow lines, respectively. The concentration of the dark state is represented by the red line. (**a**) Irreversible sequential mechanism with well separated profile peaks. (**b**) Irreversible sequential mechanism with overlapping concentration profiles. (**c**) Dead-end mechanism with separated profile peaks. (**d**) Dead-end mechanism with overlapping concentration profiles. The reaction rate coefficient for each simulation is summarized in Tab. 1. The schematic representation of irreversible sequential mechanism is shown in Fig. 1b and the dead-end mechanism is presented in Fig. 1c

To separate the electron density of pure species from a measured time series of electron density maps, methods from linear algebra are utilized such as singular value decomposition (SVD) (Schmidt et al 2003). SVD has been successfully applied to various time-resolved crystallographic data to (i) determine intermediates structures and (ii) gain information on the chemical kinetic mechanism that involves transformation between intermediates. However, SVD analysis requires expert input. In particular, the chemical kinetic mechanism needs to be estimated. Concentration profiles that are obtained by solving the coupled differential equations of the mechanism are used to obtain the pure intermediate states by using a multi-step procedure (Schmidt *et al*., 2003). This procedure (also known as projection algorithm) is (i) difficult to grasp for the non-expert, (ii) not very user friendly and (iii) ignores the direct relationship between concentration and (difference) electron density. A neural network (NN) (see Fig. 3) addresses these challenges since it allows to restore the relationship between concentration and electron density, and it is user-friendly as it does not require the user to understand the mathematics of the projection algorithm.

**Figure 3.**
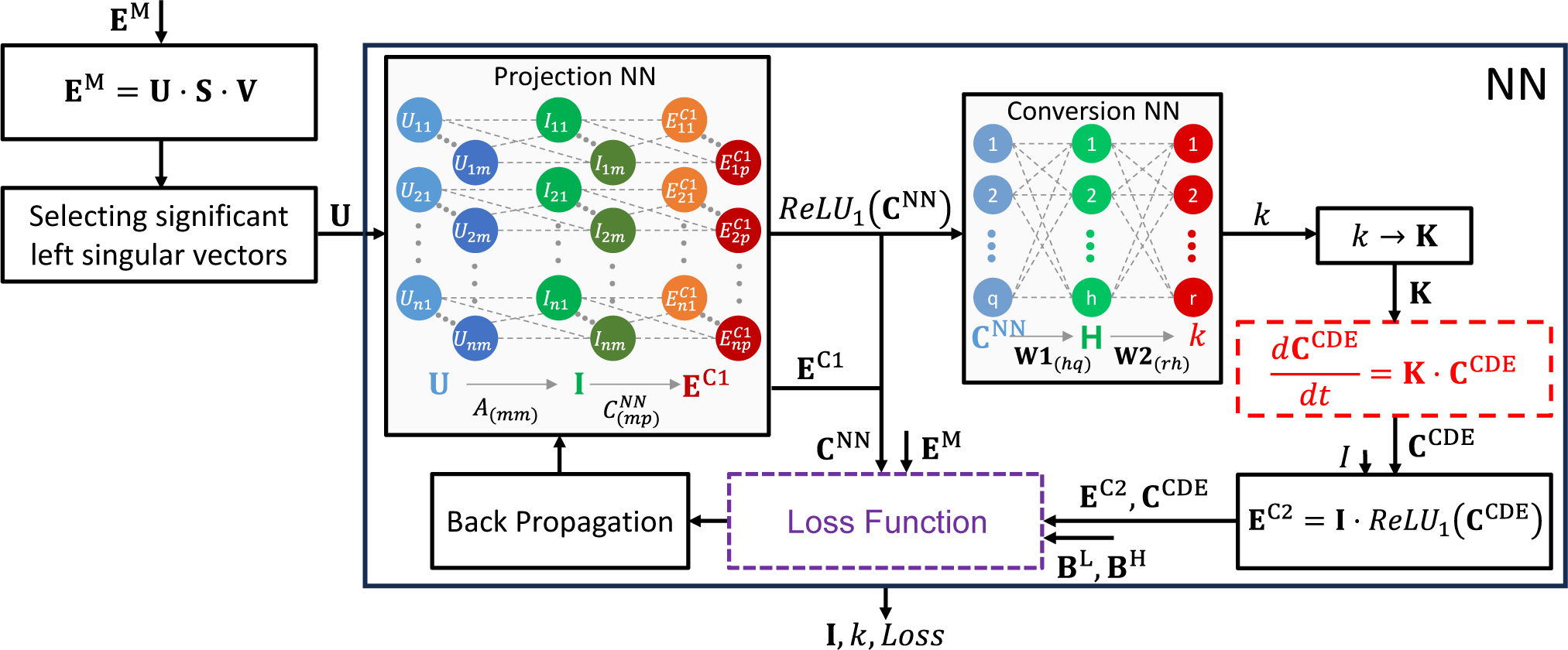
Schematic representation of the analysis method for time resolved x-ray crystallography measurements using KINNTREX. The NN consists of two sub-networks. The first network is called the projection NN. It aims to predict the input, ***E***^*M*^ (M for measured) as accurately as possible by generating time-dependent DED maps (***E***^*C1*^, C for calculated) from significant lSVs (**U**) along with the DED maps of the intermediates (***I***) and the concentrations (***C***^*NN*^). The second sub-network called conversion NN takes the ***C***^*NN*^ as input and predicts reaction rate coefficients (RRCs), ***k. C***^*NN*^ is flattened before applied to the conversion NN. After passing through both sub-networks, KINNTREX solves the differential equation governing the kinetic mechanism of the protein photocycle (red dashed box), resulting in the concentrations, **C**^CDE^. In a subsequent step, prior to the calculation of the loss function, the time dependent DED maps, ***E***^*C2*^, are predicted a second time using the DED maps of the intermediates ***I*** from the projection NN and the ***C***^*CDE*^ (eqn. 4). The loss function (purple dashed box) evaluates the discrepancies between measured and predicted time dependent DED maps as well as differences between the calculated concentrations (***C***^*NN*^ and **C**^CDE^). The user can constrain the adjustable range of the RRCs to further inform the loss function. The backpropagation procedure concludes the NN. The arrows form a loop that iterates multiple times.

NNs are artificial intelligence (AI) data processing algorithms. Inspired by the human brain, NNs are layered structures of artificial “neurons” connected to each other. Each connection is established with a different strength. The variation of the connectivity level across the network provides the ability to learn and establish a reliable output signal. To mimic the neuron in a brain, the artificial neuron is affected by a non-linear activation function. The most commonly used activation function is the so-called rectified linear unit (ReLU) (Zeiler *et al*., 2013). Neural networks are constructed with different architectures: they are called recurrent (recursive) neural networks (RNN) (Medsker & Jain, 2000), convolutional neural networks (CNN) (Gu *et al*., 2018, Fukushima, 1980), and physically informed neural networks (PINN) and its derivatives (Karniadakis *et al*., 2021, Ji & Deng, 2021, Zaverkin & Kastner, 2020, Meuwly, 2021, Westermayr & Marquetand, 2021) to name a few. Recently, X-ray crystallography has seen a growth in the use of machine learning methods and in particularly neural network algorithms for data analysis (Vollmar & Evans, 2021). Here, an NN as a member of the PINN family of networks is proposed with the goal of extracting DED maps of the intermediates and the corresponding concentration profiles from the time-resolved X-ray data alone. The NN is informed by a system of linear coupled differential equations which describe the kinetic mechanism (Steinfeld *et al*., 1999).

## 2. Background & Methods

The primary objective is to retrieve structural (conformational) changes in a protein during a reaction. In a time-resolved crystallographic experiment X-ray diffraction patterns (DP) are collected at a time Δt after the reaction has been initiated (Moffat, 1989, Srajer & Schmidt, 2017). Reflection intensities are extracted from the DP (Ren *et al*., 1999, Schmidt, 2019) from which structure factor amplitudes (|F_t_|) are calculated. Reference structure factor amplitudes (|F_ref_|) are obtained from X-ray data collected on crystals where the molecules are at rest. The structure factor phases ϕ_ref_ are derived from a well determined reference model. By subtracting the |F_ref_| from the |F_t_| difference structure factor amplitudes Δ|F|_t_ are calculated. With the help of the phases ϕ_ref_, time-dependent DED maps are obtained (Moffat, 1989, Schmidt, 2023). The goal is to extract the kinetics from a time series of DED maps. Once this is achieved, time-independent DED maps and ultimately the corresponding structures of the intermediates can be determined. This manuscript introduces a new analysis method, to recover the chemical kinetic mechanism and the DED maps of the intermediates. The new analysis utilizes a kinetically informed neural network (KINN) specifically designed to work with TRX data. This proposed NN, henceforth is named KINNTREX.

### 2.1. Neural network architecture

The laws of physics are deduced from experimental observations. These laws are described by mathematical formulations, that provide predictions for the outcome of experiments not yet conducted. In many cases, the data are so complex that an explicit prediction cannot be obtained. NNs can be employed to resolve such situations. Even though a straightforward theory or a mathematical model cannot be constructed, an NN can use existing observations to predict the outcome of experiments yet to be performed.

An elementary unit (a building block) of an NN is a perceptron (also named neuron) (Rosenblatt, 1958). A perceptron can be expressed mathematically as,

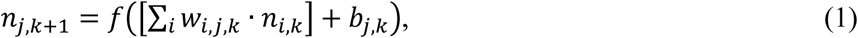

where *n*_*i,k*_ is the value in the *i*^th^ perceptron in layer *k, w*_*i,j,k*_ is the weight of the connection between perceptrons *n*_*j,k*+1_ and *n*_*i,k*_. *b*_*j,k*_ is a bias added before executing the activation function *f*. The activation function is used for changing the perceptron value in a non-linear manner, mimicking a biological neuron (Rosenblatt, 1958, Block, 1962). A common NN algorithm starts at the input layer (first layer), propagates through the following layers according to Eq. 1, and ends at the output layer. This process is called forward propagation. The output of an NN provides the essential missing information, such as the classification of input data (in cases where classification is desired). When an NN is initiated and executed for the first time, the output provides non useful relation to the input. This necessitates an iterative training process. Training involves providing inputs along with values that should be predicted by the NN (referred to as ground truth or labels). The output can be compared to the labels using a loss (cost) function, which calculates a loss value. This loss value is utilized to update the weights and biases of the NN, enabling them to be adjusted for improved performance in subsequent iterations. The updating process, known as backpropagation, employs an optimization function such as stochastic gradient decent (Jain *et al*., 1996, LeCun *et al*., 1988). Such a training procedure is called supervised training. When the training is unsupervised the ground truth is not known. This forces the NN to extract patterns from the input or classify inputs based on differences between them (Hinton *et al*., 1995, Chen *et al*., 2016). KINNTREX as introduced here is an unsupervised NN, which does, however, impose physically meaningful constraints on the data analysis.

#### 2.1.1. Data preparation for KINNTREX

Suppose a time-series of DED maps is available from a TRX experiment. Each DED map has been determined at a specific time-point Δ*t* after reaction initiation. A DED consists of a large number of data points (voxels), sampled on a three-dimensional grid which typically covers one crystallographic unit cell. Each voxel contains a difference electron density value. Structural changes are typically concentrated at a chromophore of a photoactive protein or at the active site of an enzyme where strong DED features are located. Therefore, most parts of the DED map are free of signal. This allows us to identify a region of interest (ROI) where a strong signal persists. The ROI is carved out from all difference maps in the time series (see Schmidt et al, 2003 for details). Consequently, the time-series contains much less voxels than the original DED maps.

Prior to analysis with the KINNTREX algorithm, the time-dependent ROIs are assembled and organized chronologically based on their acquisition time. Each voxel value of the ROIs is assigned to an element of a column vector. For *N* voxels in the ROI, the column vector is *N*-dimensional. This rearrangement is called flattening. The vectors are organized as a *N*×*P* matrix, **E**^M^ (E stands for difference electron density and M for measured). *P* is the number of time points.

#### 2.1.2. Singular Value Decomposition of the Matrix **E**^M^

The matrix **E**^M^ is decomposed into three matrices by singular value decomposition (SVD) (Eq. 2).

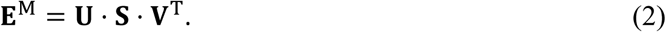

The matrix **U** (N×P) contains the left singular vectors (lSVs), and the matrix **V** (P×P) contains the right singular vectors (rSVs). The diagonal matrix, **S** (P×P) includes the singular values along its main diagonal. **V**^T^ is the transpose of matrix **V**. Matrices **U** and **V** contain significant and insignificant singular vectors, indicated by the diagonal elements of **S**. The insignificant left singular vectors are discarded, and the truncated matrix is used as input to the NN. For time-resolved X-ray data, the The selection of significant singular vectors has been discussed in detail in the literature (Schmidt *et al*., 2003).

The SVD also allows to extract relaxation rates. These values are used to constrain the NN as explained in the following subsections. The relaxation rates are obtained by fitting a sum of exponential functions to the right singular vectors (rSVs) (Henry & Hofrichter, 1992a, Schmidt *et al*., 2003). This can be formulated as follows:

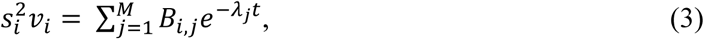

where *s*_*i*_ and *v*_*i*_ are the *i*^th^ singular value and the corresponding rSV, respectively. B_*i,j*_ is the amplitude of the *j*-th process (observed in the *i*-th rSV) which must be obtained by a fitting procedure. *λ*_*j*_ is the *j*^th^ relaxation rate, also calculated in the same fitting procedure, and *t* represents time. Note that *λ*_*j*_ is globally observed in all significant rSVs. All significant right singular vectors are fitted simultaneously according to Eq. 3, using the nonlinear least square fitting Levenberg-Marquardt algorithm (Levenberg, 1944, Marquardt, 1963). The minimum number of exponential functions that would be fitted determines the number of distinguishable processes in the reaction, and the minimum number of intermediates, M (Rajagopal *et al*., 2005, Ihee *et al*., 2005).

Once the relaxation rates are obtained, they can be used to define limits for the magnitudes of the RRCs. The RCCs are positive values, therefore, 0 is considered as the lower limit. The upper limit, however, is a multiple of the largest relaxation rate *λ*_i,_ throughout the paper. An example implementing these constraints is provided in section 3.4.

#### 2.1.3. Architecture outline

The architecture of KINNTREX is described in Fig. 3. It consists of two sub-networks, each serving a distinct purpose, as described below. Subsequently, a series of steps are implemented to solve the coupled differential equations that govern the kinetic mechanism of the protein under investigation. These steps inform the NN about the physics of the underlying protein dynamics. The calculation through the NN is repeated for multiple iterations. For each iteration the loss function is evaluated. The resulting loss value is used to monitor the progress of the calculation. The architecture is outlined by pseudo-code in Appendix A.

#### 2.1.4. First Iteration

In the initial iteration, the significant left singular vectors are loaded into the input layer of the first sub-network, referred to as the projection NN (see Fig. 3). The projection NN is a so-called partially connected, feedforward NN (Kang & Isik, 2005) as described in supplementary material section S1. In addition to the input layer, the projection NN incorporates a middle layer with perceptrons holding the calculated, time-independent DED maps of the intermediates, **I**. The calculation of the intermediates is performed using Eq. 4.

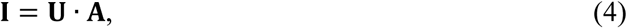

where the matrix **A** contains the weights with which the significant lSVs are multiplied. A third layer within the projection NN is dedicated to the determination of time-dependent DED maps, labeled as **E**^C1^ as calculated according to Eq. 5.

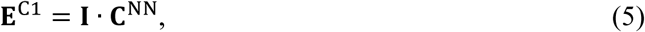

where **C**^NN^ contains the concentrations of the intermediates. The **C**^NN^ elements act as weights for the middle layer perceptron when calculating the values at the output layer. **I, C**^NN^ and **E**^C1^ are poorly predicted at this stage as only one iteration has gone through the NN.

Upon passing through the projection NN, the output, **E**^C1^, is not utilized in the subsequent sub-network. Instead, the **C**^NN^ modified by the ReLU activation function, is loaded into the second sub-network known as the conversion NN. **E**^C1^, will be used later to determine a loss value (see below). The conversion NN includes an additional hidden layer as can be seen in Fig. 3. This sub-network outputs the reaction rate coefficients (RRCs) of the chemical kinetic mechanism. The number of perceptrons in the hidden layer, *H*, is calculated according to Eq. 6,

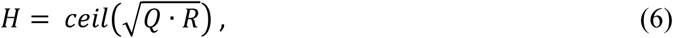

where *Q* = *M* · *P, M* is the number of intermediates, *P* is the number of time points, and *R* is the total number of RRCs within the general mechanism. *ceil*() rounds the argument to the larger closest integer. The dimensions of required matrices in Fig. 3 are shown in Tab. 3 below.

After the two sequential sub-NNs, the coupled differential equations of the chemical kinetic mechanism are solved (eq. 7):

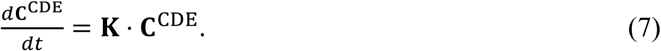

In Eq. 7, **C**^CDE^ represents the time-dependent concentrations (the concentration profiles) of the intermediates, and **K** is the coefficient matrix, assembled from the RRCs (see Appendix C). Eq. 7 is solved by diagonalizing matrix **K**. The solution of Eq. 7 is outlined in appendix B. Accordingly, the NN is informed by a general chemical kinetic mechanism containing three intermediates plus the reference state. Initial condition, **C**_ini_^CDE^ = (1, 0, 0, 0), are applied. Hence, the first intermediate concentration at *t* = 0 was set to 1 while the other two intermediates and the dark state concentrations were set to 0. The concentrations were normalized to the total number of excited molecules. All these molecules at *t*=0 can be assumed to be in their first intermediate state. If the non-excited molecules are disregarded, the concentration of the other intermediates as well as the dark state must be 0. The last step, that concludes the first iteration through the network, recalculates the time dependent DED maps, **E**^C2^, using **C**^CDE^ and **I** calculated in the middle layer of the projection NN, i.e.,

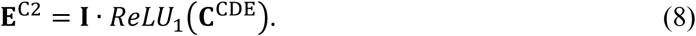

*ReLU*_1_ is similar to *ReLU* with the addition of having an upper limit of 1. Hence, values above the limit are set to the limit value. At this point **E**^C2^ and **C**^CDE^ are poorly predicted as only one iteration has been executed.

#### 2.1.5. Iterations through the NN

To monitor the progress of the KINNTREX algorithm a loss function is used. Only a portion of the resulting loss value is attributed to the comparison between the output of the NN and its input. The loss function is described in detail in section 2.2. After the loss value is determined, backpropagation is used to minimize the loss by adjusting the weights. Here we choose adaptive moment estimation (AdaM) for optimization of the weights (Kingma *et al*., 2020). The input to the optimization includes a learning rate which determines how much the weights of the NN are allowed to change. A large learning rate will result in larger change. The computation might converge faster but might converge to a less accurate prediction. For a small learning rate, the opposite is true. Thus, there is an optimal learning rate that can both converge within a reasonable time and provide a correct prediction of the desired information (Bengio, 2012). Once the loss function converged to a constant value, the intermediate DED maps, **I**, are obtained along with the RRCs of the chemical kinetic mechanism. This iterative process is both a training procedure and it provides the desired information (concentrations and DED maps of the intermediates).

### 2.2. The loss function

The loss function consists of four parts. These are, (1) the time dependent DED map loss, *L*_*E*_, (2) the reaction rate coefficient related loss, *L*_*K*_, (3) the intermediate concentration related loss, *L*_*C*_, and (4) the intermediate DED maps related loss, *L*_*I*_. The various parts are described in detail below. The total loss value is calculated as,

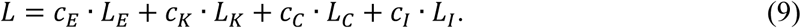

*c*_*E*_, *c*_*K*_, *c*_*C*_, and *c*_*I*_ are the user-specified amplifying factors for the *L*_*E*_, *L*_*K*_, *L*_*C*_, and *L*_*I*_ losses, respectively.

#### 2.2.1. Time dependent DED map related loss, *L*_*E*_

The first part, *L*_*E*_, calculates the differences between the input time-depended DED maps, and the two time-dependent DED maps predicted by the NN (Eq. 10):

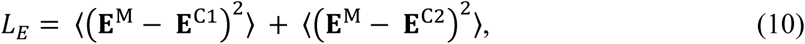

where **E**^M^ represents the input time-dependent DED maps and **E**^C1^as well as **E**^C2^ represent time-dependent DED maps as determined by the NN, respectively. The pointed brackets ⟨ ⟩ denote averaging over all matrix elements.

#### 2.2.2 Reaction rate coefficient related loss, *L*_*K*_

The second part of the loss function, *R*_*K*_, constrains the magnitudes of the RRCs (*k*_*i*_) by allowing them to be within a user-specified range. The loss function penalizes an RRC if it is found outside of the range as follows:

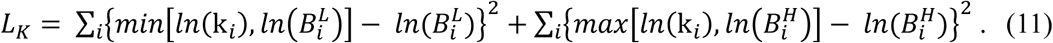

The logarithm has been used because the processes within the chemical kinetic mechanism occur at vastly different rates. This way, fast and slow processes are placed within the same order of magnetude. The *min* and *max* functions determine the minimum and maximum values between *cn*(k_*i*_) and the corresponding lower and upper limits, 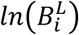 and 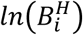, respectively. An example is shown in Tab. 2. When *k*_1_ is between the boundaries nothing is added to the loss value. When *k*_1_ is out of the range the loss value will increase. Enforcing the RRCs to stay within the boundaries will decrease *L*_*k*_. This helps the NN to converge.

#### 2.2.3 Intermediates concentration related loss, *L*_*C*_

*L*_*C*_ is determined by the difference between the time-dependent concentration profiles **C**^NN^ (from the projection NN) and **C**^CDE^ (from the solution of the coupled differential equations, Eq.7).

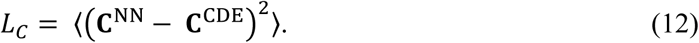

Inclusion of concentration-based loss is imperative for the method to work. This calculation forces the NN towards a self-consistent solution, as it will be shown in section 3.1.

#### 2.2.4 Intermediate DED maps related loss, *L*_*I*_

This loss consists of two parts (Eq. 13). The first part (*DED*_*ref*_) forces the reference state DED map to become zero. This part will not be calculated when the DED maps of the intermediates in the projection NN do not include the dark state. If the reference state is included the target is zero, as the reference DED map (which is featureless) is subtracted by its own prediction. The second (optional) part of this loss value demands that the DED maps of the intermediates be as different as possible from each other. This can be achieved by projecting these DED maps onto each other. The *L*_*I*_ loss value is calculated as,

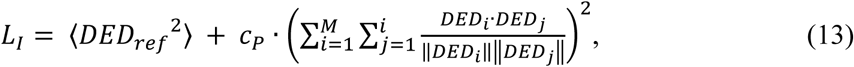

where *M* is the number of intermediates and *c*_*P*_ is a user selectable amplification factor that weighs the contribution of the projection relative to the rest of the loss value. The similarity (or dissimilarity) is calculated by the dot product between flattened vectors *DED*_*j*_ and *DED*_*i*_, where *DED*_*j*_ and *DED*_*i*_ represent time-independent DED maps of different intermediates predicted by KINNTREX. If the maps are similar, the dot product is large. This increases the loss value which the NN intends to minimize. As a result, the NN favors DED maps which are as dissimilar as possible. The doubly vertical lines (e.g. ‖*DED*_*i*_‖) denotes the *length* of the vector.

### 2.3 Implementation of KINNTREX

The NN was implemented in Python with the PyTorch package providing all the necessary tools for the implementation of the neural network. Unless indicated otherwise, the configuration of the NN was set up as follows: (i) The maximum number of iterations was set to 3x10^5^. (ii) The weights and biases were initiated with random numbers (see details in section 4.3 and supplementary material section S2). (iii) The learning rate was set to 10^-4^. (iv) The amplifying factors (*c*_*E*_, *c*_*K*_, and *c*_*I*_) (section 2.2.) were set to 1 and the factor *c*_*C*_ was set to 0.1. (v) The (optional) amplifying factor for the projection potion of the intermediate-based loss (*c*_*P*_) was set to 0. *c*_*P*_ may become useful in future applications of KINNTEX.

KINNTREX reads the ROIs extracted from a time-series of crystallographic DED maps. The ROIs are stored in ASCII format in a text file with multiple columns separated by a tab. The first column contains the voxels indexes of the ROI from the crystallographic DED maps. Subsequent columns represent the electron densities of the ROI at subsequent time points.

Apart from the time-dependent DED maps, KINNTREX also requires a model matrix to specify the general kinetic mechanism. An example of this matrix can be found in appendix C. Additional inputs include the number of intermediates and a text file with the NN parameters. The critical parameters included in the parameters file are: (i) the STD of the distribution from which the initial weights are drawn, (ii) the learning rate, (iii) the maximum number of iterations, (iv) the amplifying factors for the different loss value portions, (v) the loss tolerance, and (vi) the boundaries for the RRC ranges. If the loss value is equal to or smaller than the loss tolerance for *n* consecutive iterations, the program finishes. In case the loss tolerance is not reached, the algorithm finishes after the maximum number of iterations is reached.

KINNTREX generates the following output files, formatted as text: (i) a list of DED maps of the intermediates after each iteration, (ii) a list of RRCs after each iteration, (iii) the loss value after each iteration, and (iv) the predicted concentration profiles of the intermediates after the last iteration. The file containing the DED maps of the intermediates has *M* · *N*_*i*_ columns, where *M* is the number of intermediates and *N*_*i*_ is the number of iterations. The file contains *N* lines where *N* is the number of voxels within the selected ROI. The file containing the RRCs has *R* columns and *N*_*i*_ lines, where *R* is the number of RRCs.

The file containing the loss value has a single column with *N*_*i*_ lines. The file containing the concentrations has *M*+1 columns and *P* lines, where *P* is the number of time points. The concentrations are calculated by solving Eq. 7 using the RRCs obtained after the last iteration. In this paper, the concentration profiles typically contain about 400 time points, to ensure that the plots appear smooth over the entire time range.

In a separate module, crystallographic DED maps of the intermediates are reconstructed by mapping the resulting voxel values back to the ROI of the unit cell. An additional module calculates the residual and weighted residual values for each predicted concentration or time-dependent DED maps and the corresponding ground truth (see section 2.6). The operations performed by KINNTREX are specified by the pseudo-code listed in appendices A-C.

### 2.4. Ground Truth Simulations

The photoactive yellow protein (PYP) has been chosen as a model system in the simulations to test the algorithm. PYP is a photoreceptor found in halophilic bacteria (Meyer *et al*., 1987). PYP reacts to illumination by blue light (Sprenger *et al*., 1993, Hoff *et al*., 1994). Once it is excited it undergoes a reversible photocycle. DED maps for 21 different time points were generated by an algorithm published earlier (Schmidt *et al*., 2003). Three different intermediates and the dark/reference state were included. The structures of the intermediates were similar but not identical to intermediates I_T_, pR1, pB determined by time-resolved X-ray crystallography (Ihee *et al*., 2005, Jung *et al*., 2013). Water molecules were removed. Structure factors (SFs) for the dark state, **F**_0_, and the intermediate states, **F**_1_ to **F**_3_, were calculated using the structures of the reference state and the intermediates, respectively.

The difference structure factors of the pure intermediate states have been obtained by subtracting the structure factors of the reference state from those of the intermediate states. The three time-independent DED maps shown in Fig. 4 were calculated from these difference structure factors. Time-dependent (time-resolved) DED maps were calculated from the time-independent maps as shown below. Both simulated time-independent and time-dependent DED maps were taken as the ground truth and could be directly compared to those obtained by KINNTREX.

**Figure 4.**
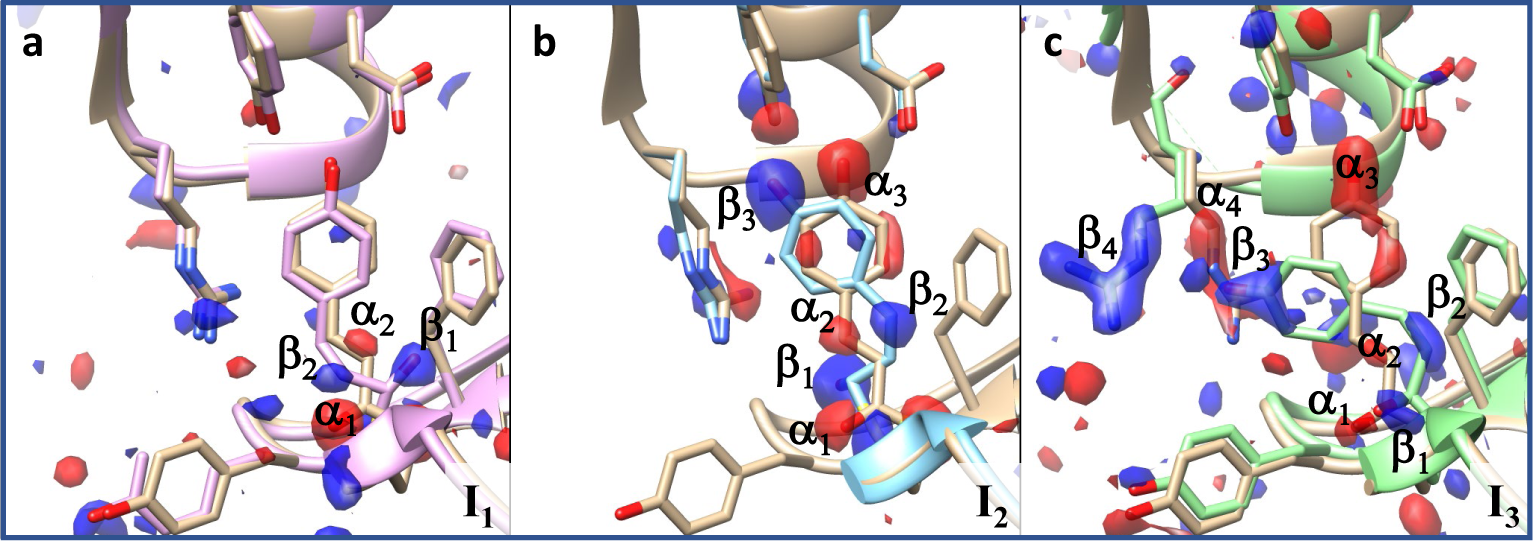
The ground truth. The simulated DED maps are overlaid onto the atomic representation of (a)the I_T_ (**b**) the pR1, and (**c**) the pB intermediate states. Negative electron density is colored red and positive colored blue. Markings label features in the DED map. ‘β’ indicate positive DED and ‘α’ indicate negative DED.

Two kinetic mechanisms have been tested: the irreversible sequential mechanism (Fig. 1b) and the dead-end mechanism (Fig. 1c). Each mechanism was simulated with two different sets of RRCs (see table 1). The concentrations of the intermediates at different time points were obtained by solving Eq. 7 with the selected RRCs as inputs and the initial condition mentioned in section 2.1.4. The concentration profiles for each simulation are shown in Fig. 2. The two sequential mechanisms are denoted as S_S_ and S_O_. The concentration profiles for S_S_ (“s” for separated) are well separated in time (Fig. 2a). For S_O_ (“o” for overlapping) the concentration profile of intermediate I_2_ is buried within that of intermediate I_1_ (Fig. 2b). Two other concentration profiles, DE_S_ as well as DE_O_, were generated representing the dead-end mechanism. Again, in DE_S_ the intermediates concentrations are separated (Fig. 2c). For DE_O_, the I_3_ concentration profile is fully buried within the I_2_ concentration profile (Fig. 2d).

**Table 1.** Reaction rate coefficients (RRCs) with units 1/s used to generate the concentration profiles in Fig. 2. The mechanisms and RRCs are also used to simulate ground truth data for the NN. Based on the mechanism employed and the extent of overlap of the concentrations, the simulations are referred to as S_S_ and S_O_ for sequential separated and sequential overlapped, and DE_S_ and DE_O_ for dead-end separated and dead-end overlapped, respectively.

| RRC | S<sub>s</sub> | S<sub>o</sub> | DE<sub>s</sub> | DE<sub>o</sub> |
| --- | --- | --- | --- | --- |
| $k_1$ | 40,000 | 2,000 | 9,500 | 15,000 |
| $k_3$ | 1,000 | 3,000 | 330 | 2,000 |
| $k_4$ | - | - | 400 | 100 |
| $k_5$ | 100 | 900 | - | - |
| $k_{-3}$ | - | - | 210 | 2,000 |

**Table 2.**
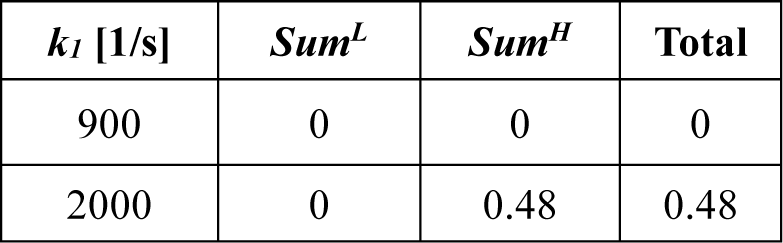
Illustration of the loss value *L*_K_. A case where one of the RRCs, *k*_*1*_, is within the boundaries, and another where *k*_*1*_ is outside the boundary is shown. *Sum*^*L*^ and *Sum*^*H*^ equal the left and right summation terms in Eq. 11, respectively. The *L*_*k*_ for *k*_*1*_ is calculated in the fourth column. The lower and upper limits 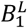 and 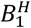 are set close to 0 (zero excluded, because ln 0 is not defined) and 1000 1/s, respectively.

### 2.5 Time-dependent difference maps

The time-dependent SFs were calculated from the SFs of the intermediates **F1, F2, F3** and the dark state **F**_**0**_ as follows,

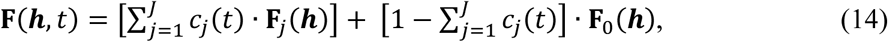

*c*_*j*_(*t*) is the concentration of the *j* intermediate at time *t*. Noise was added to the amplitudes of the SFs. The noise was estimated from the experimental standard deviations found in a Laue data set collected from a PYP dark state crystal (for details, see Schmidt et al., 2003). Difference SF amplitudes were calculated by subtracting reference state (dark state) SF amplitudes with noise from the noisy time-dependent SF amplitudes. A weight factor calculating the weighted difference structure factor amplitudes has been implemented for the purpose of reducing very large differences resulting from experimental noise (Ren *et al*., 2001). With the weighted amplitudes and the phases calculated from the dark state structural model, time-dependent DED maps were obtained by Fourier transformation using the CCP4 program ‘FFT’ (Winn *et al*., 2011). DED features were pronounced near the chromophore region. Accordingly, this region was chosen as the ROI. The ROIs from different time points were assembled into matrix **E**^M^ and decomposed by SVD. The significant lSVs (columns of matrix **U**) were taken as the input for KINNTREX (Fig. 3). Tab. 3 shows the dimensions of the matrices used by KINNTREX in this study.

**Table 3.** Dimensions of matrices and vectors in KINNTREX required with the simulated data. Upper: Dimensions of matrices **E, U, A** and **C** in the projection NN of KINNTREX (see Fig. 3). In the simulations, *M* = 3 processes are predicted from *P* = 21 time-points measured during a reaction and *N* = 2291 voxels are in the ROI of each DED map. For each matrix, the first subscript denotes the number of rows and the second the number of columns. Lower: Vectors and matrices in the conversion NN. The vector **C**^NN^ is determined by assembling the rows of matrix **C**^NN^, from the projection NN, into a vector. The dimension *H* in **W1** and **W2** must be larger than *R*. Matrices **E**^M^, **E**^C1^ and **E**^C2^ as well as **C**^NN^ and **C**^CDE^ are comparable, respectively, and contribute to the loss value.

|  | Projection NN |  |  |  |  |
| --- | --- | --- | --- | --- | --- |
| | $\mathbf{E}_{(n,p)}^{\text{M}}$ | $\mathbf{U}_{(n,m)}$ | $\mathbf{A}_{(m,m)}$ | $\mathbf{I}_{(n,m)}$ | $\mathbf{C}_{(m,p)}^{\text{NN}}$ |
| rows | $N = 2291$ | $N = 2291$ | $M = 3$ | $N = 2291$ | $M = 3$ |
| columns | $P = 21$ | $M = 3$ | $M = 3$ | $M = 3$ | $P = 21$ |
|  | Conversion NN |  |  |  |  |
| | $^*\mathbf{C}^{\text{NN}}$ | $\mathbf{W1}_{(h,q)}$ | $\mathbf{H}$ | $\mathbf{W2}_{(r,h)}$ | $\mathbf{k}$ |
| rows | $Q = M \cdot P = 63$ | $H = 25^*$ | $H = 25$ | $R = 10$ | $R = 10$ |
| columns | 1 | $Q = 63$ | 1 | $H = 25^{**}$ | 1 |

\* Converted from $\mathbf{C}_{(m,p)}^{\text{NN}}$ by subsequently lining up the three rows
\*\* $H$ must be greater than $R$ (see Eq. 6)

### 2.6. Accuracies of KINNTREX

The loss value *L*_*E*_ indicates how close the output time-dependent DED maps are to the input data. The accuracies of the concentrations and those of the time-independent DED maps of the intermediates cannot be estimated, since in an experiment these ground truths are unknown. Optimization of the critical parameters (such as learning rate, maximum number of iterations, boundaries of RRC ranges, etc.) can be done by using simulated data. In addition, suitable parameters can be found by observing the reproducibility for multiple runs of the NN (see results in section 4.3). Once the network parameters are found, KINNTREX is primed to work with DED maps of the measured protein of interest.

To compare the concentrations with the ground truth, the weighted residual (*R*_*w*_) was chosen. The *R*_*w*_ was calculated by using the predicted concentrations and the ground truth (simulated) concentrations as follows:

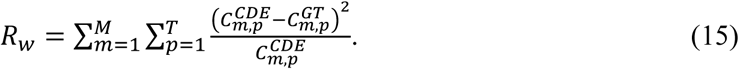

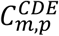 and 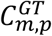 are the predicted and ground truth concentrations for the intermediate m, at the time point p, respectively. The first summation in Eq. 15 includes the reference state. Zero values of 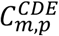 were ignored to avoid making *R*_*w*_ infinite.

For experimental data, a residual (*R*_*s*_) can be calculated for the measured and predicted time-dependent DED maps. The *R*_*s*_ is similar to the DED map loss value portion, *L*_*E*_. To calculate *R*_*s*_ Eq. 16 is used.

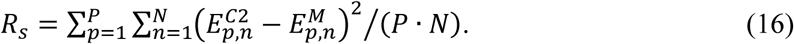

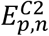 and 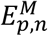 are the predicted and measured time-dependent DED maps, respectively. *N* is the voxel count and *P* is the number of time points.

To evaluate the reliability of the results, KINNTREX must be executed at least ten times. Each attempt is executed with different random initial values for the weights and biases. The predicted concentrations are extracted from the last iteration at each execution. Once the concentrations are obtained, *R*_*w*_ and *R*_*s*_ values are calculated using Eq. 15 and Eq. 16, respectively. Following the residuals calculations, the values are assembled into histograms. The goodness of fit estimation for the data in this paper is discussed below (section 4.3) and elaborated in section S2 of the supplementary material.

## 3. Results

The performance of KINNTREX was tested with a variety of experimentally realistic scenarios, simulating protein transformation through various chemical kinetic mechanisms. In the first scenario, the NN was tested with data generated by the irreversible sequential mechanism S_S_. In this case, concentration profiles of the intermediates exhibit well-separated maximum values (Fig. 2a). For that, KINNTREX is given the following a-priori information. (i) The number of intermediates obtained after the initial SVD analysis. (ii) A general mechanism containing 3 intermediates with 10 rate coefficients (Fig. 1a). (iii) At *t* = 0, only the first Intermediate is populated, all other concentrations are zero. (iv) All weights and biases of the NN were initialized by drawing random values from a Gaussian distribution with 0 mean and STD of 0.02. No additional restrictions on rate coefficients were imposed.

In the second scenario, a more challenging irreversible sequential mechanism with overlapping intermediate concentration profiles (S_O_) was tested (see Fig. 2b). To overcome this challenge, the NN was subjected to constraints by limiting values of the RRCs to a certain range (Eq. 11). Relaxation rates obtained from the right singular vectors were used to determine the limits (see section 2). The maximum values of all RRCs were set to multiples of the fastest relaxation rate, and the minimum values were set to essentially zero. See Eq. 11 and Tab. 2 for examples of their effect on the loss function. This constraint was added to the a-priori information provided for the former scenario. In fact, this information was provided to KINNTREX while executed for all the scenarios.

In Scenario 3 data simulated from a dead-end mechanism were investigated. This mechanism includes an equilibrium between two intermediates. Such an equilibrium is not present in the irreversible sequential mechanism. The mechanism is depicted in Fig. 1c. The dead-end mechanism introduces another complication, as individual concentration profiles can exhibit multiple transients. Consequently, the multi-featured concentration profile can be misinterpreted as a relaxation of two different intermediates instead of the transition of a single intermediate towards two other intermediates. As an additional challenge to KINNTREX, a severe overlap of two concentration profiles was also introduced in this scenario.

In scenario 4 a concentration profile with a hardly visible second transient is generated (Fig. 2d). This scenario may constitute a significant challenge, as the second (weak) transient may not be recoverable by KINNTREX.

### 3.1. Scenario 1, the S_S_ mechanism

To test the first scenario (S_S_), KINNTREX was executed with no restrictions imposed upon all ten RRCs of the general mechanism (Fig. 1a) (for practical reasons the low and high limits of all RRCs were set to essentially 0 and 10^10^ 1/s, respectively). Fig. 5 summarizes the prediction along with the ground truth for the S_S_ simulation. The predicted RRCs (Fig. 5b - solid lines) closely align with the ground-truth curves (Fig. 5b - dashed line). Additionally, the predicted concentrations (Fig. 5c - solid lines) exhibit a strong correlation with the ground truth values (Fig. 5c - circles). The loss value converged close to 7·10^-5^ (Fig. 5a)

**Figure 5.**
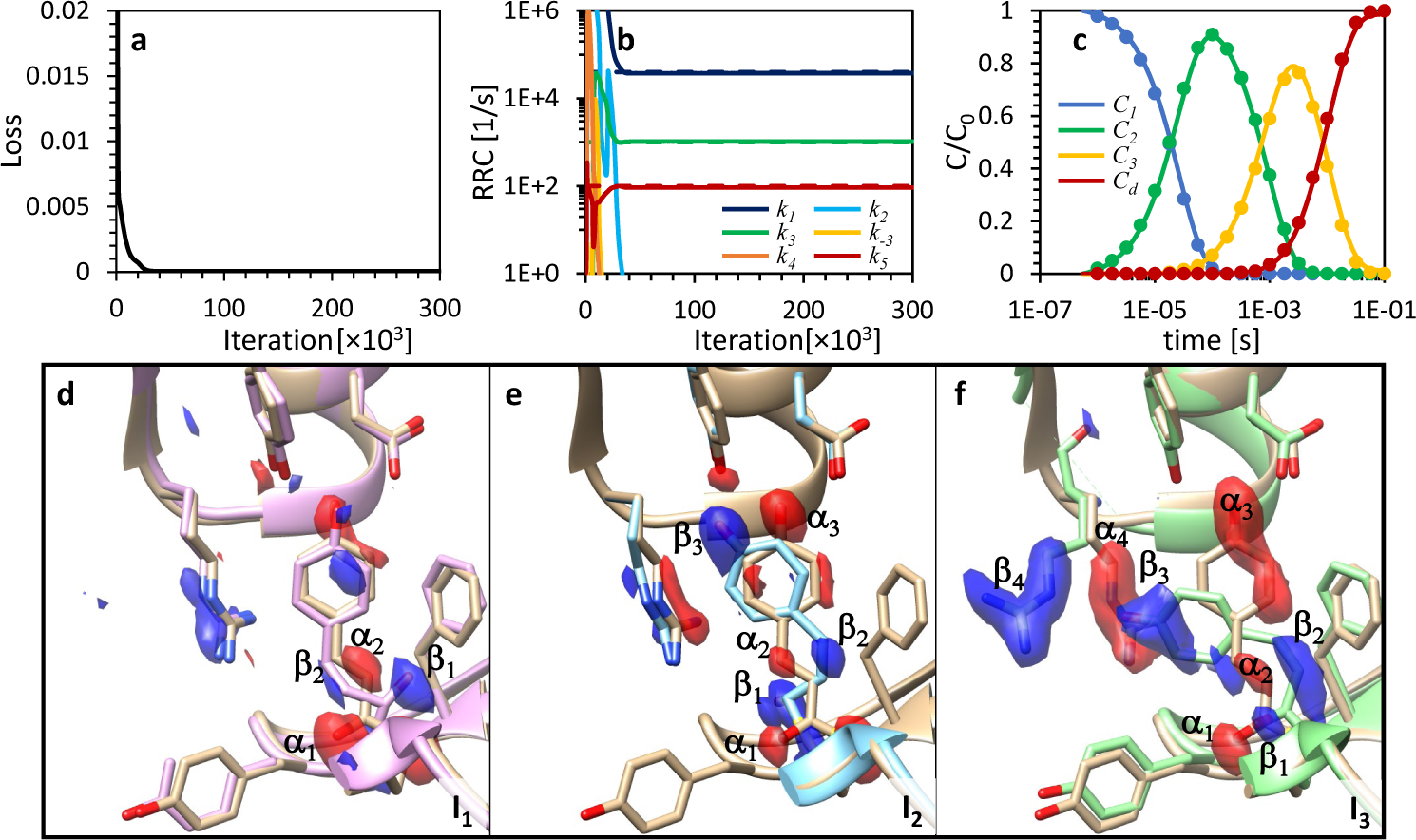
RRCs, concentration profiles and DED maps of the intermediates as obtained by KINNTREX from time-dependent DED maps generated by the S_S_ mechanism. (**a**) Loss value vs iteration number. (**b**) predicted RRCs vs iteration number (solid lines) along with the ground truth value (dashed line). (**c**) Temporal evolution of the relative concentrations of the intermediates at the last iteration (solid lines) along with ground truth (circles). The concentration profiles of intermediates *I*_*1*_ to *I*_*3*_ are shown by blue, green and yellow lines respectively, whereas that of the reference state *I*_*d*_ is shown in red. (**d**) DED maps of the intermediates (**I**_**1**_, **I**_**2**_, and **I**_**3**_ as marked in the figure) overlaid on top of their ground-truth atomic structure as well as the reference atomic structure (light brown). Negative electron density is colored red and positive colored blue.

In addition to fully recovering the chemically kinetic mechanism and RRCs, the DED maps of the intermediates (Fig. 5d) are very close to the ground truth DED maps, as shown in Fig. 4. Both sets of DED maps (from Fig. 4, and form Fig. 5, panels d, e, and f) display negative and positive electron densities at corresponding positions.

This result is truly remarkable, as KINNTREX has retrieved the kinetic mechanism along with the intermediate DED maps without any underlying assumptions guiding the analysis. This accomplishment can be attributed to two pivotal factors: First, the concentration values are intricately embedded in the time evolution of the DED maps. The NN capitalizes on its ability to recognize patterns. The pattern recognition is performed by comparing two sets of time-dependent DED maps in the NN: one from the projection NN (**E**^C1^) and the other calculated by Eq. 8, following the solution of the coupled differential equations (**E**^C2^). Both sets of DED maps are compared against the input DED maps (**E**^M^) and their differences are minimized iteratively. Second, informing the NN by a chemical kinetic mechanism, through coupled differential equations, forces the NN to converge towards the correct answer.

### 3.2. Scenario 2, the S_O_ mechanism

The second scenario involves a more complex irreversible sequential mechanism where two of the concentration profiles overlap (Fig. 2b). concentration profile of I_2_ is buried in that of I_1_. When the NN is executed with the same constraints as applied in scenario 1, the prediction was unsatisfactory, as evident in Fig. 6a and Fig. 6d. None of the predicted RRC follow the ground truth at any iteration (Fig. 6a). Apart from the dark state concentration, none of the intermediate concentrations follow the ground truth. The concentration for the first intermediate is 0 for all time points (Fig. 6d). To improve the prediction, the RRCs were restricted to a range of reasonable values. The upper limit (3×10^4^ 1/s) was set about ten times the highest relaxation rate value 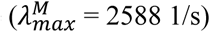, the lower limit is essentially zero.

**Figure 6.**
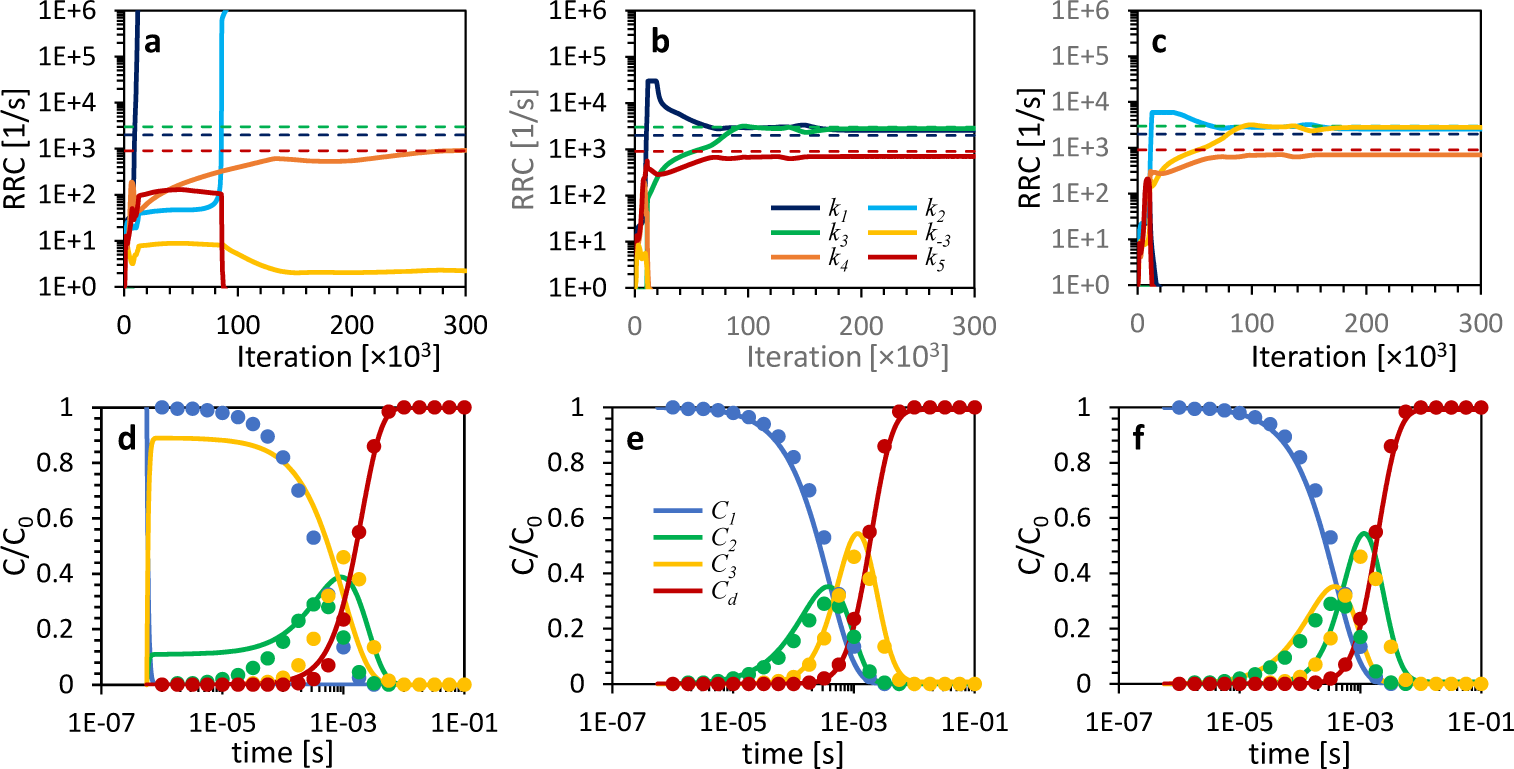
KINNTREX retrieval of the RRCs and the concentration profiles of the intermediates from time-dependent DED maps, constrained by 3 different sets of RRC range limits. (**a**-**c**) RRCs vs iteration number for the *S*_*O*_ simulation with lower and upper range limits of (**a**) 0 and 10^10^, (**b**) 0 and 3×10^4^, and (**c**) 0 and 6000, respectively. The ranges were the same for all the 10 RRCs. (**d**-**f**) Relative concentration profiles of the intermediates for the various simulations as predicted by the NN (solid lines) along with ground truth (circles). Each bottom panel graph (d, e, and f) is calculated with the RRCs extracted at the last iteration as presented in the panels a, b, and c, respectively.

Fig. 6b and Fig. 6e show a significant improvement of the prediction for the RRCs and the concentration profiles, respectively. The RRCs and the concentration profiles (solid lines) coincide well with the ground truth (circles). Further tightening the ranges of the RRCs by a factor of 5 only led to incremental improvement as illustrated in Fig. 6c and Fig. 6f. In Fig. 6f the concentration profiles for I_2_ (green) and I_3_ (yellow) are interchanged. Inspection of the mechanism (Fig. 1a) reveals that the reaction path from I_1_ to I_3_ and I_2_ is a valid representation of the same sequential mechanism as used for the ground truth.

Using ten adjustable RRCs, the DED maps of the intermediates and the kinetic mechanisms can be predicted correctly without a prior knowledge of the exact mechanism. Hence, unlike in the SVD analysis (Schmidt *et al*., 2003), where a specific mechanism had to be imposed, KINNTREX eliminates the need for such an assumption.

### 3.3. Scenario 3, the dead-end mechanism

The dead-end mechanism adds another level of complexity due to reversible process between intermediates I_2_ and I_3_. This mechanism is depicted in Fig. 1c. DE_O_ exhibits overlapping concentration profiles, as indicated in Fig. 7c by the green and yellow points.

**Figure 7.**
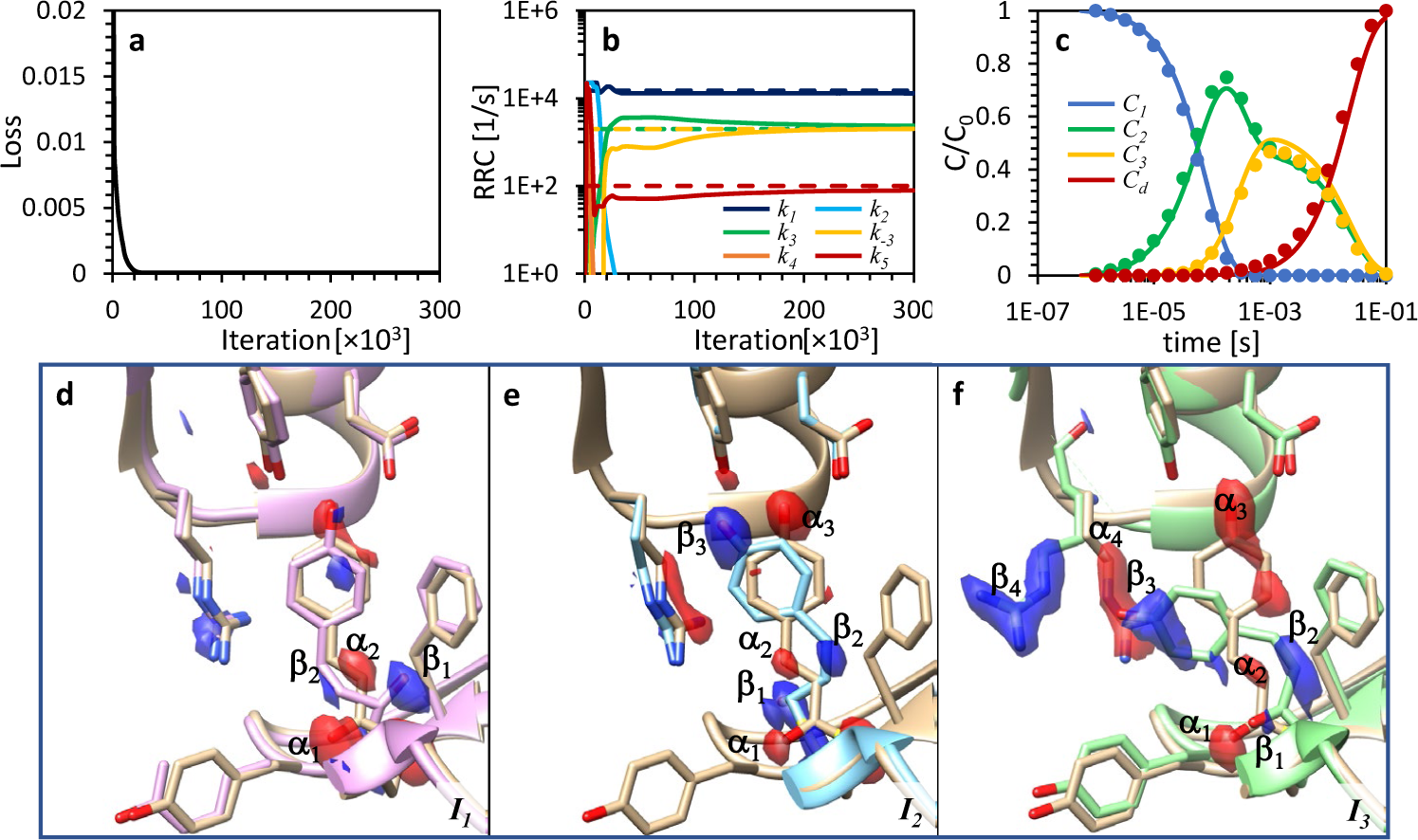
RRCs, concentration profiles and DED maps of the intermediates as predicted by KINNTREX from time-dependent DED maps for the DE_O_ mechanism. The loss value calculation includes the comparison between the two calculated concentration profiles, **C**^NN^ and **C**^CDE^ (i.e., *L*_*c*_). (**a**) Loss value vs iteration number. (**b**) predicted RRCs vs iteration number (solid lines) along with the ground truth values (dashed line). (**c**) Temporal evolution of the relative concentrations of the intermediates as predicted by the NN at the last iteration (solid lines) along with ground truth (circles). The concentration profiles of intermediates I_1_ to I_3_ are shown by blue, green and yellow lines respectively, where that of the reference state I_d_ is shown in red. (**d**) DED maps of the intermediates (**I**_**1**_, **I**_**2**_, and **I**_**3**_ as marked in the figure) overlaid on top of their atomic structure as well as the reference atomic structure (light brown). Negative electron density is colored red and positive colored blue. The RRC boundaries were set to essentially 0 and 1.5×10^5^ 1/s for the lower and upper limits, respectively.

The adjustable range limits for all ten RRCs were set to essentially 0 and 1.5×10^5^ 1/s for the lower and upper limits, respectively. The upper limit is about 10 times the value of the largest identifiable relaxation rate 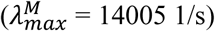.

The lightly restricted NN retrieved both the RRCs (Fig. 7b) and the concentrations (Fig. 7c) as evident by the overlap of the predicted concentration profiles with those of the ground truth (dashed lines in Fig. 7b and circles in Fig. 7c). Again, the loosely restricted NN with no assumptions on the mechanism produced results remarkably close to the ground truth, despite the complications imposed by the reversible process.

### 3.4. Scenario 4, the dead-end mechanism – with a small second transient along the concentration profile

Scenario 4 considers data simulated with the DE_S_ mechanism. In this scenario two complications were introduced. (i) A reversible process between two intermediates, I_2_ and I_3_. (ii) The second transient in the concentration profile of I_2_ buried when substantial noise is added to the simulated data. The results from KINNTREX are summarized in Fig. 8. The first attempt uses a constrained NN with lower and upper limits set to essentially 0 and 9500 1/s for all ten RRCs (Fig. 8a and Fig. 8d). The upper limit equals to the sum of the relaxation rates obtained from the SVD analysis. This is more restricted than the loose limit of 10 times the highest relaxation rate used for scenarios 2 and 3. As can be seen from Fig. 8a a dead-end-like mechanism was not recovered. However, the four RRCs with most-significant magnitude (*k*_*1*_, *k*_*3*_, *k*_*4*_, and *k*_*5*_) produced concentration profiles very close to the ground truth. The second transient feature of the I_2_ concentration profile was not recovered (Fig. 8d, green line and arrow). In addition, concentrations predicted for intermediate I_3_ are slightly higher than the ground truth (Fig. 8d, yellow line).

**Figure 8.**
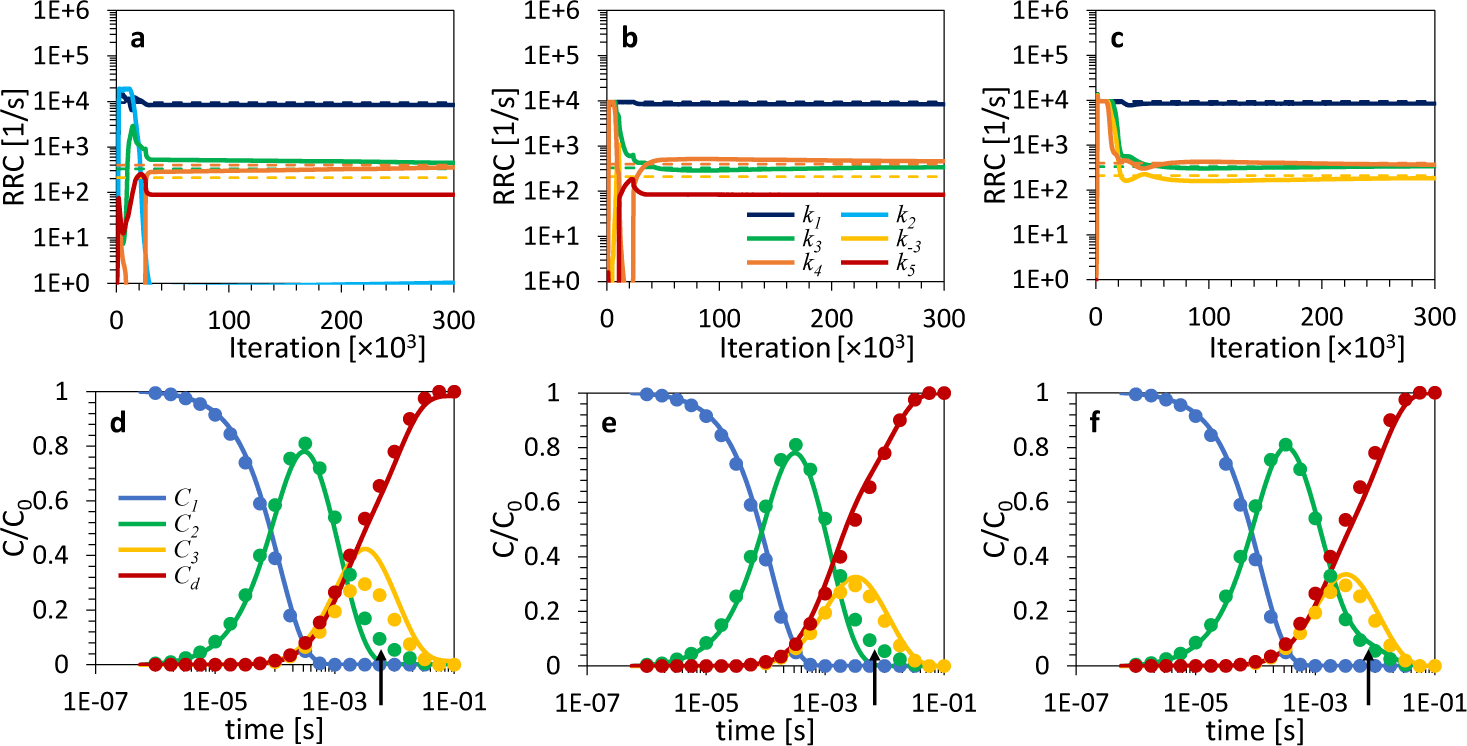
KINNTREX retrieval of the RRCs and the concentration profiles of the intermediates from time-dependent DED maps constrained by 3 different sets of RRC range limits. (**a**-**c**) RRCs vs iteration number for the DE_S_ simulation with lower and upper limits of (**a**) essentially 0 and 9500 for all ten RRCs, (**b**) essentially 0 and 9500 for 5 RRCs (*k*_*1*_, *k*_*3*_, *k*_*4*_, *k*_*5*_, and *k*_*-3*_), and (**c**) 84 and 9500 for 4 RRCs (*k*_*1*_, *k*_*3*_, *k*_*4*_, and *k*_*-3*_), respectively. (**d**-**f**) Relative concentration profiles of the intermediates for the various simulations as predicted by the NN (solid lines) along with ground truth (circles). Each bottom panel graph is calculated with the RRCs extracted at the last iteration as presented in the panel above. The upper limits for the RRCs are in all cases (**a**-**c**) equal the sum of the observed relaxation rates while the lower limit for (**c**) equals the smallest of the relaxation rates. The arrows indicate the weak second transient in the concentration profile of *I*_*2*_.

To recover more accurate concentrations, additional constraints have been imposed. Five out of the ten RRCs were forced to zero (Fig. 8b and Fig. 8e). Upper and lower limits of the other five RRC (*k*_*1*_, *k*_*3*_, *k*_*4*_, *k*_*5*_, and *k*_*-3*_) were set close to 0 and 9500 1/s, respectively. Still, the obtained concentration profile for intermediate I_2_ did not reveal the second transient (see arrow in Fig. 8e), but the concentration profile for I_3_ has improved.

In a third attempt six RRCs were set to zero, with the remaining four RRCs (*k*_*1*_, *k*_*3*_, *k*_*4*_, and *k*_*-3*_) constrained between 84 and 9500 1/s (Fig. 8c and Fig. 8f), where the lower bound was set to the magnitude of the lowest relaxation rate obtained from the SVD analysis (see section 2.1.2). The four adjustable RRCs corresponded to those of the dead-end mechanism depicted in Fig. 1c. This time, the predicted concentration profile for the I_2_ intermediate reveals the second transient (arrow in Fig. 8f) and all the other predicted concentration profiles agree well with the ground truth. However, in this case, KINNTREX was informed by the underlying mechanism. Unless the mechanism can be identified by other means, small admixtures of I_2_ into I_3_ are unavoidable. In practice, these admixtures may be so small that they remain undetectable in the DED maps, and do not affect the recovery of the DED maps of the intermediates in a significant manner.

## 4. Discussion

The RRCs in the KINNTREX algorithm required constraints to provide accurate results. It can be argued that for the simplest scenario involving the irreversible sequential mechanism S_S_, no constraints are needed. For all other scenarios, adding constraints on the *k*s was necessary for accurate results. This is demonstrated below.

### 4.1. KINNTREX overcoming protein kinetics challenges

Several challenges were introduced into the kinetic mechanisms by simulation of different scenarios. These added challenges include: (i) overlapping concentration profiles of the intermediates, (ii) reversible processes, and (iii) transients within a concentration profile which are difficult to detect. While the first two complications were tackled by constraining the RRC ranges relatively relaxed (about 10 times the value of the largest relaxation rate), the third complication required a more restrictive approach. Informing KINNTEX with a dead-end mechanism, albeit, with unknown reaction rates, was demonstrated in section 3.4. The dead-end mechanism was informed by forcing six of the ten RRCs to zero. The other four RRCs, which participate in the dead-end mechanism, were constrained to a range. Constraining KINNTREX by informing it with the mechanism is discussed in the following subsection.

#### 4.1.1. Effect of small second transients on KINNTREX

As shown for scenario 4 in section 3, the concentration profile for the I_2_ intermediate contains a second small transient (Fig. 8). Even restricted constraints could not retrieve the concentration profile accurately. Accordingly, several kinetic mechanisms have been selected and tested. A comparison between the attempt to retrieve data by informing KINNTREX with the dead-end and sequential mechanisms is presented in Fig. 8 and Fig. S4, respectively. The sequential mechanism forced seven RRCs to zero with the remaining three (*k*_*1*_, *k*_*3*_, and *k*_*5*_) constrained between 84 and 9500 1/s. Informing the KINNTREX with the dead-end mechanism resulted in acceptable predictions. Fig. 8c shows overlap between the retrieved RRCs (solid lines) and the ground truth (dashed lines). Fig. 8f shows that the predicted concentrations agree closely with the ground truth. However, informing KINNTREX with the irreversible sequential mechanism (Fig 1b) resulted in a failed prediction. As evident in Fig. S4a only two of the three RRC in the sequential mechanism were found to be significantly larger than zero. Furthermore, a total mismatch between the retrieved concentration profile (Fig. S4b solid line) and ground truth (Fig. S4b circles) is evident. The loss values calculated for both informed mechanisms can be compared for choosing the appropriate kinetic mechanism. A lower loss value indicates better accuracy. KINNTREX converges to a loss value of 6.4×10^-5^ for the dead-end mechanism and 12.9×10^-5^ for the sequential mechanism. These values agree with the conclusion from Fig. 8 and Fig. S4 that the dead-end mechanism is the favored one. Furthermore, *R*_*w*_ for the dead-end mechanism is 0.1 and that for the sequential mechanism is 496 (see section 2.5 for the estimation of *R*_*w*_) which clearly favors the dead-end mechanism.

### 4.2. Degenerate chemical kinetic mechanisms

Typically (except for the S_S_ mechanism), the KINNTREX may retrieves a mechanism that differs from the underlying mechanism used in the simulation. One such example is observed in the results presented in Fig. 7b and Fig. 7c. In this case, KINNTREX predicts a mechanism that resembles the reversible sequential mechanism shown in Fig. 9b, while the data were simulated with the dead-end mechanism depicted in Fig. 1c. In such a case, the concentration profiles of the intermediates and the corresponding DED maps are essentially identical for both mechanisms. This degeneracy is impossible to break with data collected at a single temperature. Potentially, by varying the temperature as suggested earlier (Schmidt *et al*., 2010, Schmidt *et al*., 2013), the degeneracy can be lifted. It is very appealing to implement changes to KINNTREX that can analyze this type of 5D crystallographic data.

**Figure 9.**
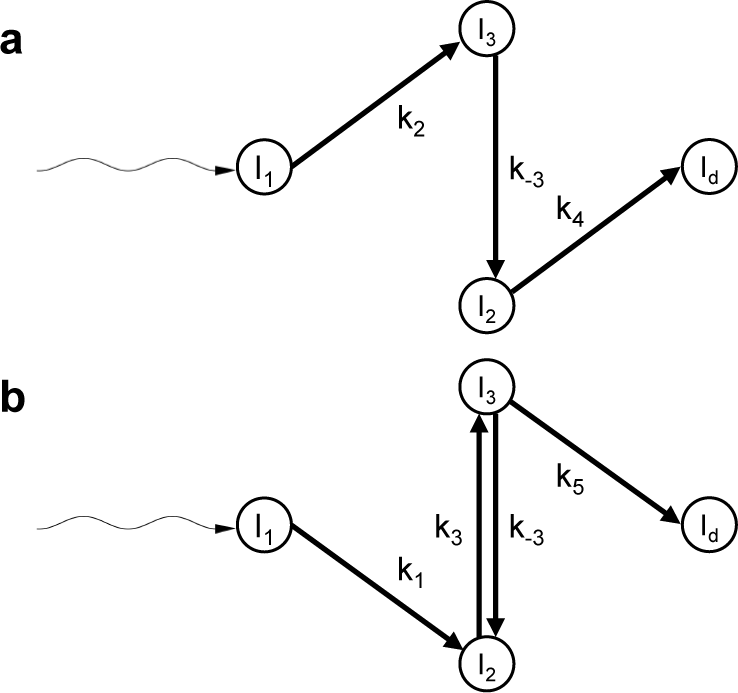
Chemical kinetic mechanisms with three intermediates for the (a) irreversible sequential mechanism, and (b) sequential mechanism with reversable process. The wavy arrow represents the light excitation of the protein. Circles represent the intermediate states and straight arrows represent transformation between intermediates. The labels on the arrows represent the reaction rate coefficients (RRCs).

### 4.3. Testing the accuracy of the predictions

There is some concern that initiating the NN weights and biases may affect the prediction significantly. To assess this, the NN is randomly initialized with different values for its weights and biases while keeping the rest of the parameters the same. KINNTREX was executed 100 times for each scenario. To speed up the analysis, a computer cluster was utilized with parallel executions. Using the cluster reduced the computational time by two orders of magnitude.

The distributions of weighted residuals (R_w_, Eq. 15) of the concentrations are narrow (Fig. 10) no matter how the weights and biases are initiated. The *R*_w_ peak values show that the more complex the data become the higher the peak values (Fig. 10, blue, green, red and yellow lines, respectively, for the different scenarios), but still the distributions remain narrow (Fig. 10). In conclusion, weights and biases do not affect the prediction.

**Figure 10.**
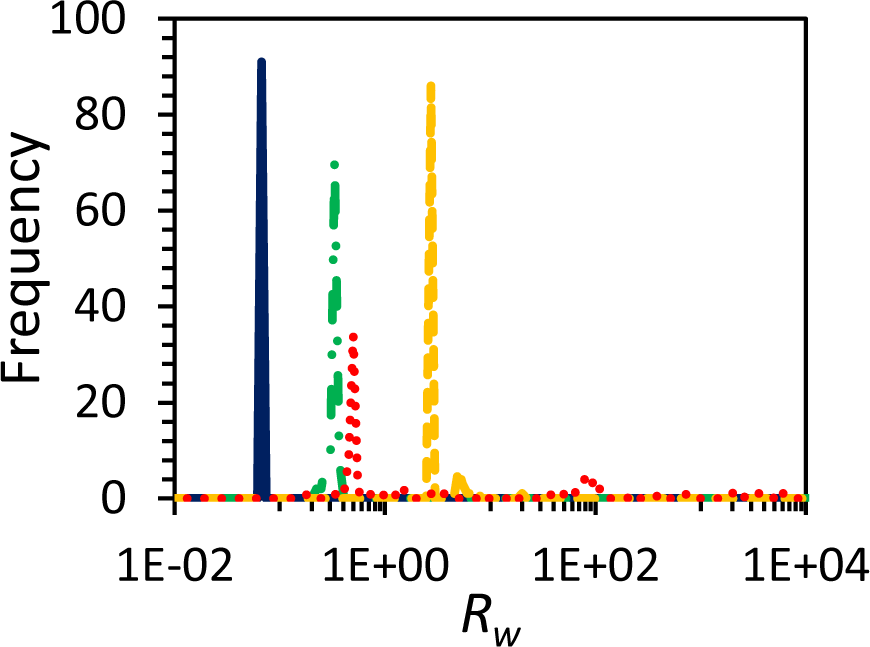
Goodness of fit distribution for the S_S_ (blue solid line), S_O_ (green dash-dotted line), DE_S_ (yellow dashed line), and DE_O_ (red dotted line) scenarios described in section 3. Weighted residual (*R*_*w*_) histogram calculated between ground truth and predicted concentrations (Eq. 15). The distributions were assembled from 100 executions of KINNTREX for each scenario with different initial weights for each execution (see section 2.5 for goodness of fit description).

In a real-world experiment, the ground truth for the concentration profiles is not known. Therefore, the residual, *R*_*s*_, as introduced in section 2.5 is used. The distributions of the *R*_s_ values are also very narrow. Even more, the peak values are all around ∼3.4·10^-5^ (electrons/Å^3^) independent of the complexity of the simulated data. The *R*_s_ peak values are comparable to the noise in the input data, which was extracted by reconstructing the data matrix **E**^**M**^ without the significant singular vectors and values (Eq. 2). For all scenarios KINNTREX predicted correct time-dependent DED maps. If the residual is significantly larger than the noise level, KINNTREX’s predictions are most likely inaccurate; see supplementary material, section S2 for an example.

## 5. Conclusions

Successes of NNs have been demonstrated recently by the popular AlphaFold2 (Jumper *et al*., 2021) and GPT3 algorithms (Vaswani *et al*., 2017). However, GPT3 and AlphaFold2 are trained with a enormous amount of data. In contrast, KINNTREX needs only the data from a single experiment and physically and chemically reasonable constraints. Such physics informed neural networks have been proven to be data-efficient as discussed in the literature (Raissi *et al*., 2019).

## 6. Outlook

This manuscript presents first groundbreaking results employing a NN to analyze time-resolved X-ray data. The NN successfully extracts kinetic mechanisms, time-dependent concentrations, and DED maps of intermediate states for several challenging scenarios. More details can be added to improve the performance of KINNTREX, such as forcing the DED maps of the intermediates to be as different from each other as possible (see part 2.2.4 in the loss function). For example, more hidden layers can be added to the conversion NN shown in Fig. 3. The linear coupled differential equation solver (shown by the red dashed box in Fig. 3), can be replaced by a non-linear type. In this way, processes which include higher-order reactions or processes involving diffusion of substrate or ligand (Malla *et al*., 2023, Olmos *et al*., 2018, Pandey *et al*., 2021), could be analyzed as well.

Now, KINNTREX needs to be applied to experimental data. For the benefit of a wide user community, KINNTREX must be equipped with an intuitive GUI. In addition, KINNTREX needs to be extended for reconstruction of the entire unit cell from the DED maps. This requires applying the symmetry operations and periodic boundary conditions, similar to what has been done previously for the application of the SVD to TRX data (Schmidt *et al*., 2003). The reconstructed content of the unit cell will then be subject to a Fourier-transform to obtain difference structure factors (DSFs). By extrapolating these DSFs,, conventional electron density maps can be obtained that enables modelling of a structure for each intermediate (Schmidt, 2023). The implementation will be a formidable challenge for scientists in the years to come.

With some modifications, the proposed method can also be applied to different types of experimental data, such as those obtained by time-resolved small or wide-angle X-ray scattering (Cammarata *et al*., 2008, Cho *et al*., 2010, Dmitri & Michel, 2003, Putnam *et al*., 2007), or data obtained by time-resolved absorption spectroscopy. Analyzing time-resolved absorption spectra could be beneficial as measurements are performed on less costly, more accessible instruments. Additionally, such results can be complementary to those obtained from time-resolved X-ray crystallography (Zimanyi, 2004, Nagle *et al*., 1995).

## Acknowledgements

This work was enabled by the NSF Science and Technology Center Biology with XFELs (BioXFEL), NSF-STC 1231306.

## Appendix A: Pseudo-code for KINNTREX

This pseudo-code describes the KINNTREX algorithm with the architecture shown in Fig. 3 (see section 2.1).

1. Decompose matrix **E**^M^ by SVD and estimate the number of significant singular values.
2. Estimate the relaxation times from the signification right singular vectors (see section 2.1.2).
3. Construct ‘Prediction NN’ with an input layer, **E**^M^, a middle layer for intermediate DED maps, **I**, and output layer for the recalculation of time dependent DED maps, **E**^C1^ (see sections S1). as **C**^NN^.
4. 1. Initiate weights matrices between input and middle layers (**A)** and between middle and output layers **(C**^NN^).
5. Construct ‘Conversion NN’ with a hidden layer and initiate the weights and biases.
6. Set constraints for the RRCs using the relaxation times from step 3.
7. **For** iteration, *c* ≤ *N*_*i*_
8. Calculate **E**^C1^ along with **I** and **C**^NN^ using forward propagation of the ‘Projection NN’.
9. Calculate the RRCs vector,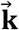, using forward propagation of the ‘Conversion NN’ with **C**^NN^ from step 8.
10. Calculate Matrix **K** using RRC vector according to equations C3.4 and C3,5.
11. Obtain **C**^CDE^ by solving Eq. 7 from matrix **K** and the initial condition **C**^ini^ using the pseudo code from Appendix B.
12. Calculate **E**^C2^ using Eq. 8 with **I** from 9 and **C**^CDE^ from step 12.
13. Calculate loss function according to Eq. 9.
14. Calculate backpropagation using the optimization method AdaM.
15. After every *m*^*th*^ iteration step, save **I**, RRCs, and the loss value.
16. **End loop**

## Appendix B: Pseudo-Code for Solving differential equations by diagonalization of the K matrix

1. Diagonalize **K** and find the eigenvalues *λ* along with corresponding eigenvectors, **Λ**.
2. **C**alculate 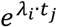 where *λ*_*i*_ is the *i*^th^ eigenvalue and **t**_*j*_ is the *j*^th^ time point.
3. Calculate **Y**(*t* = 0) = **Λ**^−1^ · **C**^ini^, where **C**^ini^ denotes the initial concentrations and **Y**(**t** = 0) corresponding value in the diagonalized space.
4. Calculate 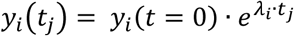
5. Finally, calculate **C**^CDE^(*t*) = **S** · **Y**(**t**).

## Appendix C: Calculating the K Matrix from RRCs

The dimensions of the matrix **K** are *D x D*, with. *n*_*I*_ + 1 where *n*_*I*_ is the number of intermediates. The addition of 1 account for the dark state. To calculate the coefficients matrix, **K**, RRCs vector,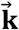, is multiplied by the so-called ‘model matrix’ **M**,

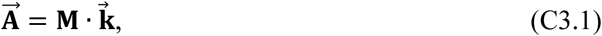

where the vector 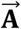 is later reshaped into the **K** matrix. The vertical dimension, *V*, of the model matrix is calculated as,

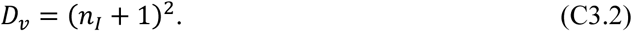

The horizontal length, *H*, equals to the dimension of the RRC vector which is also a function of the number of intermediate states and is calculated as,

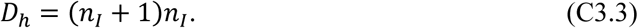

In the specific model employed here it is assumed that intermediate state 1 does not relax directly to the dark state. Vice versa, once the molecules arrive in the dark state, they cannot return to intermediate state 1 without illumination. Consequently, the number D_h_ is smaller by two compared to the number calculated from Eq. C3.3. If *n*_*I*_ equals 3 the number of relevant RRCs is equal to 10 (and not 12 as expected from Eq. C3.3). D_h_ = 10 is used here throughout (Fig. 1a). The model matrix used for the simulations on PYP in this paper is given by,

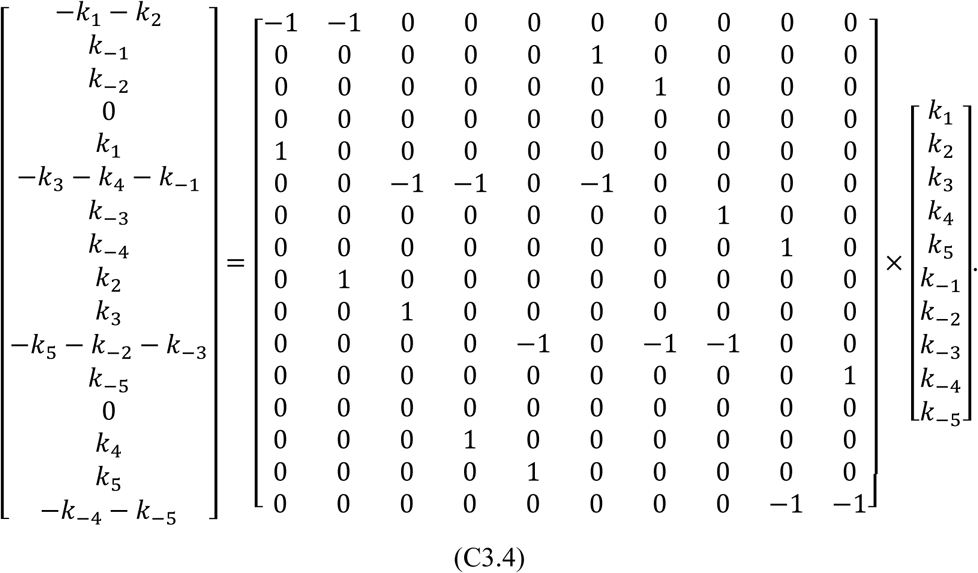

Finally, the coefficient matrix **K** is obtained by reshaping vector 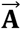 (lefthand side of Eq. C3.4),

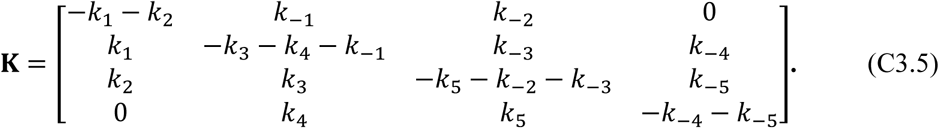

Note that the column sums of matrix **K** are zero. This guarantees the sum of all concentration remain constant for all time point. In some cases, certain rate coefficients are constrained to be zero.

## Appendix D: Converting from concentrations to RRCs using KINNTREX

The concentration profiles of the intermediates are calculated twice for each iteration. First time they are extracted from the weights in the projection NN. Second time, the concentrations are obtained by solving the coupled differential equations of the chemical kinetic mechanism (Eq. 7) by diagonalization. The RRCs are contained in the coefficient matrix **K**. To retrieve the RRCs, a simple fully connected neural network is used. This network has an input layer which is the flattened concentration vector determined from the projection NN, a hidden layer, and an output layer containing the resulting RRC values. The hidden layer perceptrons hold the weighted sum of the input values followed by an application of an activation function Leaky-ReLU, (Maas *et al*., 2013). The output layer perceptrons hold the weighted sum of the hidden layer perceptron values, but this time no activation function is applied. The sub-network is part of a larger neural network and thus its output is not compared to ground truth values but used as an input to the calculation of the coupled differential equations. The reason for using Leaky-ReLU is to avoid zero values.

## Supplementary Material

**Figure S1.**
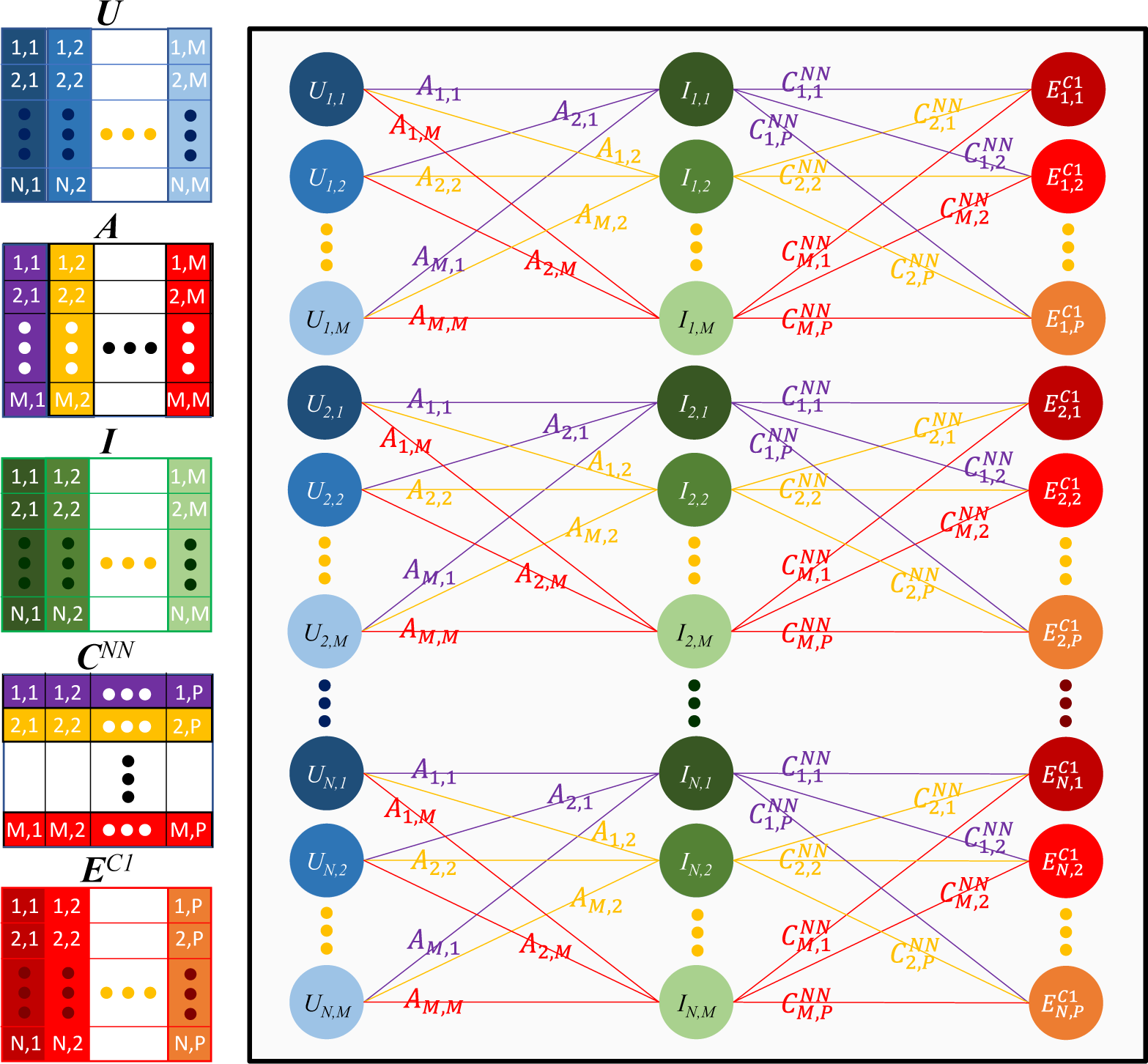
Schematic representation of the Projection NN sub-network. On the right, the network is represented by neurons (circles) and connections (lines) with weights assigned to the connections as indicated by text placed on the connection lines. The first layer **U** is the input layer (blue circle). Time-independent DED maps of the intermediates **I** are obtained in the second layer (green circles), and the red or orange circles represent the time dependent DED maps as an output of the NN. The dimensions of each matrix within the NN is presented in the left side of the figure. The columns of **U** contain the significant left singular vectors. **A** is the projection matrix. **A** is a square matrix containing weights with which the input layer is multiplied. **C**^NN^ is a weights matrix whose rows contain the concentration profiles of the intermediates. Matrix **I** is multiplied with the weights matrix **C**^NN^ to obtain the output matrix (the time-dependent DED maps). Different colors within the **C**^NN^ matrix indicate concentration profiles of different intermediates. Tab, 2 in the main text provides an example for matrix dimensions.

### S1. Partially connected neural networks

The input is an *N*×*M* matrix in Fig. S1 labeled **U**. This matrix contains the left singular vectors (lSV). *N* is the number of grid points in the region of interest (ROI) carved out from the time-dependent DED maps. *M* is the number of significant lSVs from the SVD analysis, which is also the number of distinguishable intermediates. The matrix **A** can be associated with the projection matrix of the SVD analysis (Abraham & Chain, 1940, Henry & Hofrichter, 1992b) or the projection algorithm (Schmidt *et al*., 2003). Each entry in matrix **A** determines what fraction of an lSV belongs to a particular intermediate. Multiplying **U** by **A** results in an *N* × *M* matrix containing the DED values of the intermediate states. The weights used for the calculation of the output layer are represented by matrix **C**^NN^ which is an *M* × *P* array, where *P* is the number of time points. These weights are equivalent to the time-dependent concentrations of the intermediate (concentration profiles). The resulting time-dependent DED maps are located in the output layer in a *N* × *P* dimensional matrix. The projection NN is partially connected. This scheme can still be considered an NN because of the non-linear activation function (ReLU) applied to **C**^NN^. The matrix **C**^NN^ is used as the input in the following sub-NN (conversion NN).

**Figure S2.**
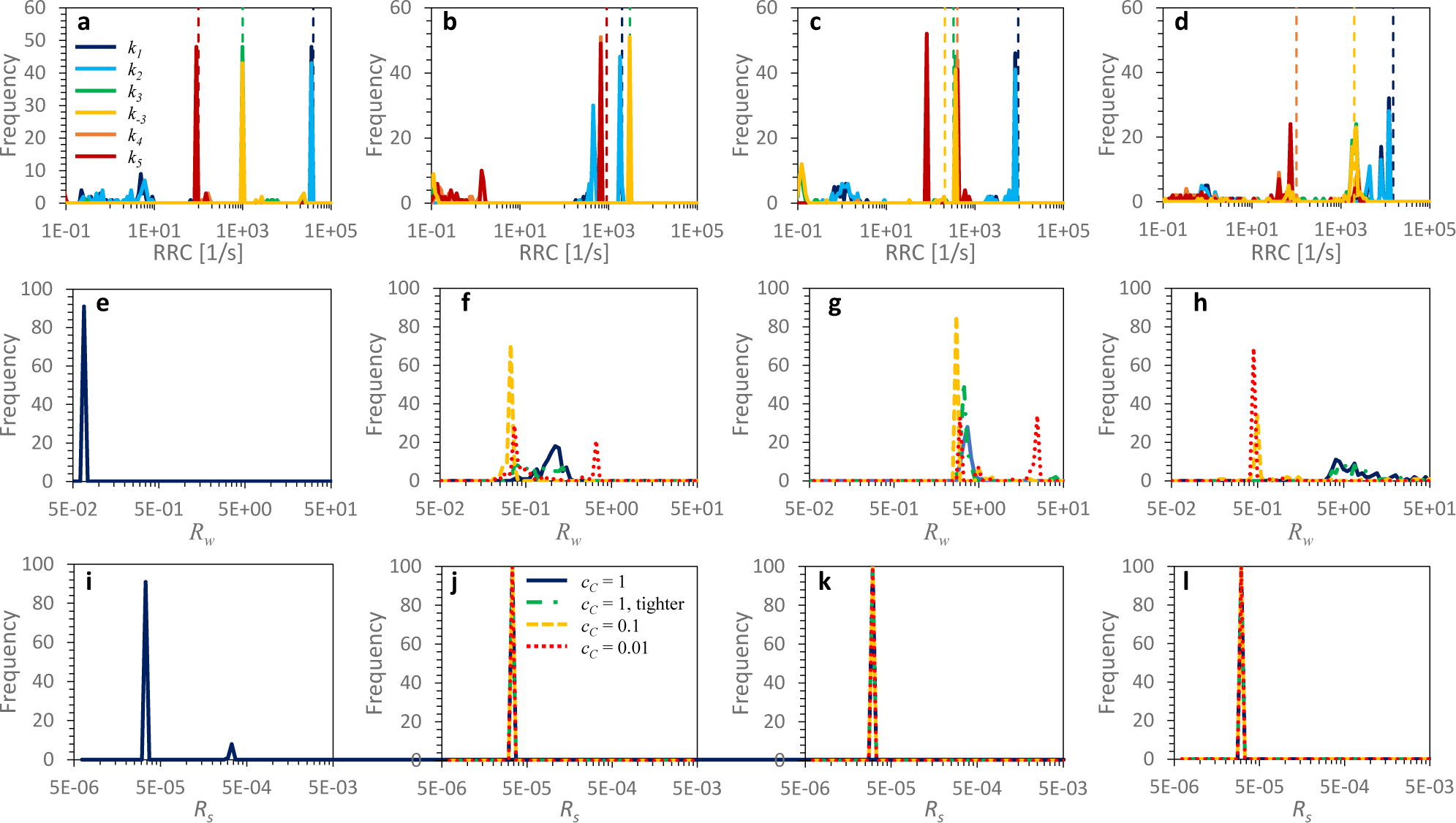
Influence of the amplifying factor *c*_C_ for the concentrations in the loss function on the RRCs and the residuals *R*_w_ and *R*_S_. Distributions of *k*_*1*_(dark blue line), *k*_*2*_ (light blue line), *k*_*3*_ (green line), *k*_*-3*_ (yellow line), *k*_*4*_ (orange line), and *k*_*5*_ (red line) for predictions of (**a**) the S_S_, (**b**) the S_O_, (**c**) the DE_S_, and (**d**) the DE_O_ scenarios. Dashed lines represent the ground truth. The amplifying factor for the concentration (*c*_*C*_) equals 0.1. (**e**-**h, middle row**): distributions of *R*_*w*_ for the different scenarios. Each panel corresponds to the panel above. The different conditions include *c*_*C*_ = 1 (dark blue line), *c*_*C*_ = 1 with tighter constraints on the RRC ranges (green dashed dotted line), *c*_*C*_ = 0.1 (yellow dashed line), and *c*_*C*_ = 0.01 (red dotted line). (**i**-**l, bottom row**) Distributions of the residual *R*_*s*_ for the different scenarios. Each panel corresponds to the panels above. The colors of the different *c*_*C*_ in the bottom row of panels are similar to those in the middle row. The distributions were assembled from 100 independent executions of KINNTREX for each scenario with a given *c*_*C*_.

### S2. Effect of the NN random initiation

In KINNTREX the matrices **A, W1, W2, C**^**NN**^, and the biases are initiated with random numbers drawn from a Gaussian distribution with 0 mean and 0.02 standard deviation. The initiation may affect the outcome and thus must be tested. The RRCs were chosen to evaluate the tests along with the figures of merits *R*_*w*_ (Eq. 15) and *R*_*s*_ (Eq. 16). *R*_*w*_ is the weighted residual, calculated between the predicted and ground truth concentrations of the intermediates. *R*_*s*_ is the residual calculated using the input and predicted time-dependent DED maps. In this study, 100 independent executions of the NN were performed for each scenario S_S_, S_O_, DE_S_ and DE_O_, respectively. In addition, the amplification factors for the concentrations (*c*_*C*_, see Eq. 9) were varied from 1.0, 0.1, 0.01 to 0.0. Figure S2 shows the results. The distributions of the RRCs predicted for the S_S_ scenario (Fig. S2a – solid lines) follow the ground truth (dashed lines) closely. The distributions are narrow indicating that the predictions of the RRCs are reproducible. Hence, the initial values of the weights and biases have little effect on the results. Fig. S2e shows that the peak of the weighted residual *R*_*w*_ is narrowly distributed around 0.07. This value is the lowest among all the scenarios. This result agrees with the precise predictions of the RRCs shown in Fig. S2a. Fig. S2i shows essentially that the same small residual value of 3.48·10^-5^ is obtained from all attempts. For the scenarios S_O_ and DE_O_ the predictions of the RRCs are presented in Fig. S2b and S2d, respectively. In both cases the predicted RRCs follow the ground truth closely and the distributions are narrow. The distributions of the *R*_*w*_ (Fig. S2f and Fig. S2h) are narrow when the *c*_*C*_ is smaller or equal to 0.1 and the peak positions of the *R*_*w*_ are 0.34 and 0.5, respectively for the two scenarios. For the *c*_*C*_ > 0.1 the *R*_*w*_ peaks are above 0.45 and 4, respectively for the different scenarios and are more spread. Wider distribution indicates a lack of reproducibility. For *c*_c_ = 0.1 we get the best results with our type of data for all scenarios. For KINNTREX the initiation of other weights and biases does not play an important role since the distributions are small. Since the R_w_ is not accessible to real data, an optimum *c*_c_ may be determined with simulated data using the protein under investigation and a suitable ROI as input to KINNTREX.

**Table S1.** Reaction rate coefficients and total loss values calculated at the last iteration of the NN scheme for the D_O_ simulation. The table compares between different realizations of the same sample with 8 adjustable reaction rate coefficients and added loss calculation for equating **C**^NN^ to **C**^CDE^.

| | $c_c = 1$ | $c_c = 0.1$ | $c_c = 0.01$ | $c_c = 0$ | $c_c = 1, \text{ Tight}$ | Ground Truth |
| --- | --- | --- | --- | --- | --- | --- |
| $k_1$ | 12088 | 12088 | 12088 | 12088 | 12088 | 15000 |
| $k_2$ | 12088 | 12088 | 12088 | 12088 | 12088 | 0 |
| $k_3$ | 3641 | 2208 | 1998 | 4024 | 3641 | 2000 |
| $k_4$ | 49 | 74 | 81 | 49 | 49 | 100 |
| $k_5$ | 45 | 74 | 81 | 49 | 45 | 0 |
| $k_{-1}$ | 0 | 0 | 0 | 0 | 0 | 0 |
| $k_{-2}$ | 0 | 0 | 0 | 0 | 0 | 0 |
| $k_{-3}$ | 3641 | 2208 | 2208 | 4024 | 3641 | 2000 |
| $k_4$ | 1 | 0 | 0 | 2 | 1 | 0 |
| $k_5$ | 1 | 0 | 0 | 2 | 1 | 0 |
| $R_w$ | 4 | 0.5 | 0.45 | 735 | 5 | |
| $R_s$ | $3.3 \cdot 10^{-5}$ | $3.3 \cdot 10^{-5}$ | $3.3 \cdot 10^{-5}$ | $9 \cdot 10^{-5}$ | $3.3 \cdot 10^{-5}$ | Noise=$3.2 \cdot 10^{-5}$ |

As indicated by Tab. S1 for the DE_O_ scenario the only RRCs that should have values different from zero are *k*_*1*_, *k*_*3*_, *k*_*-3*_, and *k*_*4*_. However, predictions have resulted in other RRCs with much larger values such as k_2_ equals 12088 and k_5_ has values equal and higher than 45. Tab. S1 shows results from a statistical analysis of 100 executions of KINNTREX for each case. Hence, in some instances the kinetic mechanism follows a sequence shown in scheme S1a while others follow a sequence presented in scheme S1b. Obviously, both schemes represent the same mechanism (Fig 1a). KINNTREX cannot distinguish between these two schemes and randomly chooses either the one or the other. In both cases the concentration profiles as well as the DED maps of the intermediates are predicted correctly.

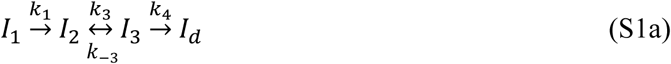

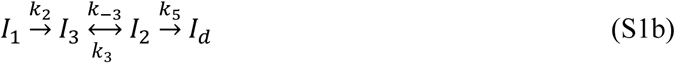

KINNTREX was initiated with weights and biases values drawn from a uniform distribution set between -1 and 1. This attempt was tested using scenario *S*_*O*_. The results were not different from the ones using Gaussian distribution with a standard deviation of 0.02 centered around 0.

### S3. Ignoring the comparison between C^NN^ and C^CDE^ in the loss function by setting *c*_*c*_ = 0

**Figure S3.**
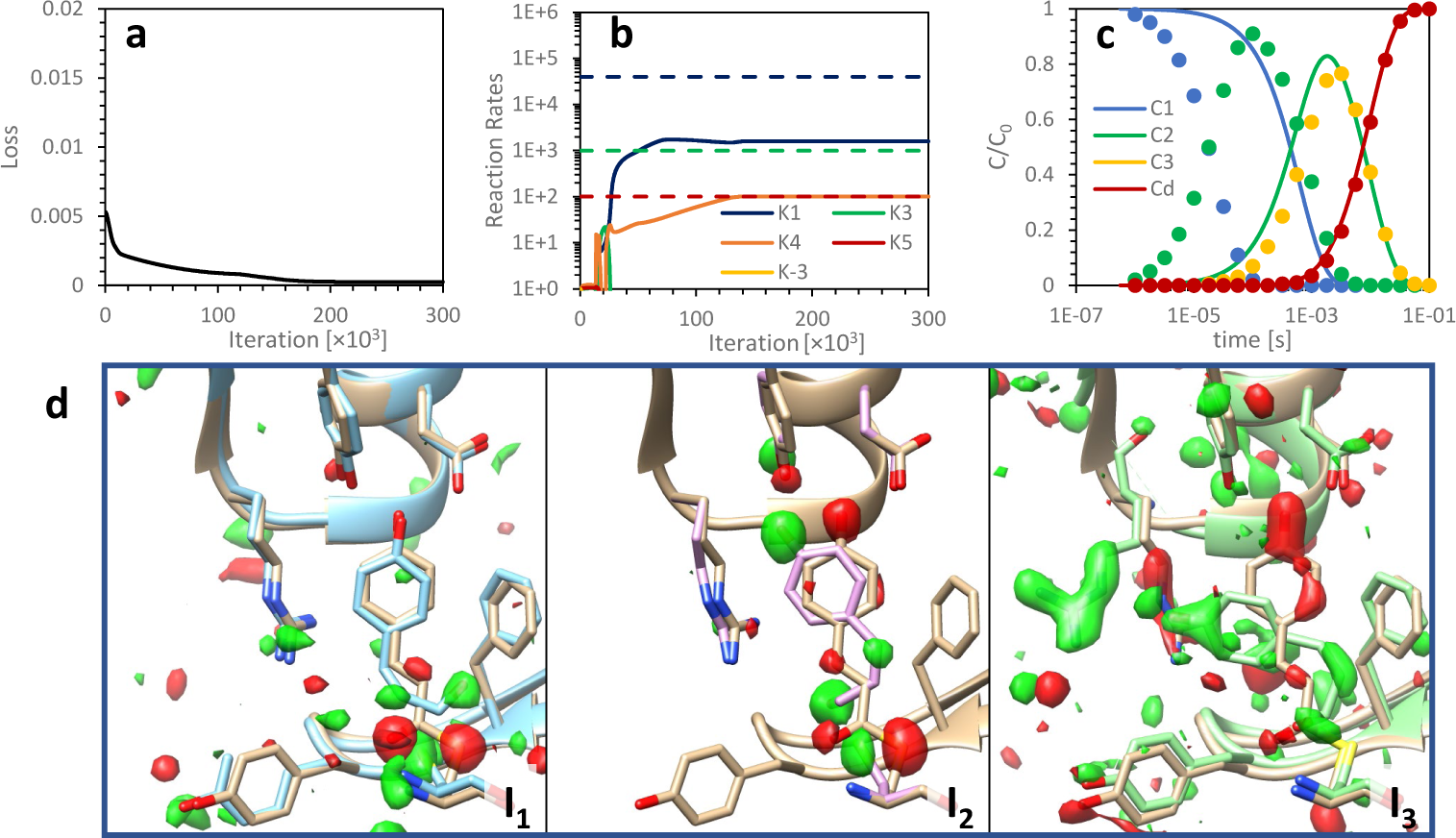
RRCs, concentration profiles and DED maps of the intermediates as predicted by the NN from time dependent DED maps for the S_S_ mechanism. The loss value calculation exclude comparison between two calculated concentration profiles, ***C***^*NN*^ and ***C***^*CDE*^. (**a**) Loss value vs iteration number. (**b**) predicted RRCs vs iteration number along with the ground truth value (dashed line). (**c**) Temporal evolution of the relative concentrations of the intermediates as predicted by the NN at the last iteration (solid lines) along with ground truth (circles). C1 to C3, Cd in the figure legend represent the concentrations at intermediates I_1_ to I_3_ and I_d_, respectively, where I_d_ is the reference (dark) state. (**d**) DED maps of the intermediates (**I**_**1**_, **I**_**2**_, and **I**_**3**_ as marked in the figure) overlaid on top of their atomic structure as well as the reference atomic structure (light brown). Negative electron density is colored red and positive colored green. The RRC boundaries were set to 0 and 10^10^ for the lower and upper limits, respectively

When the *c*_*C*_ equals zero, KINNTREX is executed without the information on the kinetic mechanism. KINNTREX is unable to predict sensible concentration profiles and DED maps (Tab. S1 and Fig. S3). *R*_*w*_ is 4 orders of magnitude larger than for the other cases, and *R*_*s*_ is about 3 times higher than the noise level while the other cases have values very close to the noise level. KINNTREX predicts 2 RRC when 3 are required (Fig. S3b). Fig. S3c shows a total mismatch between the predicted and the ground truth concentration profiles. The comparison between ***C***^*NN*^ and ***C***^*CDE*^ is imperative. Already the smallest weight *c*_*C*_ = 0.01 has been sufficient for a meaningful prediction.

### S4. Effect of setting individual RRCs to zero

Setting individual RRCs to zero during the execution of KINNTREX is equivalent to enforcing a specific candidate chemical, kinetic mechanism from the general mechanism. KINNTREX is then informed with the candidate mechanism with relevant RRCs and unknown magnitudes. Fig 8 shows the result when NN is informed with a correct candidate, in this case the dead-end mechanism. Fig. S4 presents the prediction of KINNTREX when it was informed with the sequential mechanism, but the simulated input data correspond to the dead-end mechanism. The prediction is very poor. Fig. S4a shows that only two of the three RRC in the sequential mechanism were predicted to be different from zero. Fig. S4b shows the mismatch between the predicted and ground truth concentration profiles (*R*_*w*_ = 496). Since with experimental data *R*_*w*_ cannot be evaluated, the *R*_*s*_ needs to be calculated and compared when various candidate mechanisms are employed. The candidate with the lowest *R*_*s*_ needs to be selected. In general, candidates can be selected to explicitly test and refine several scenarios, but in the general case, this is not necessary.

**Figure S4.**
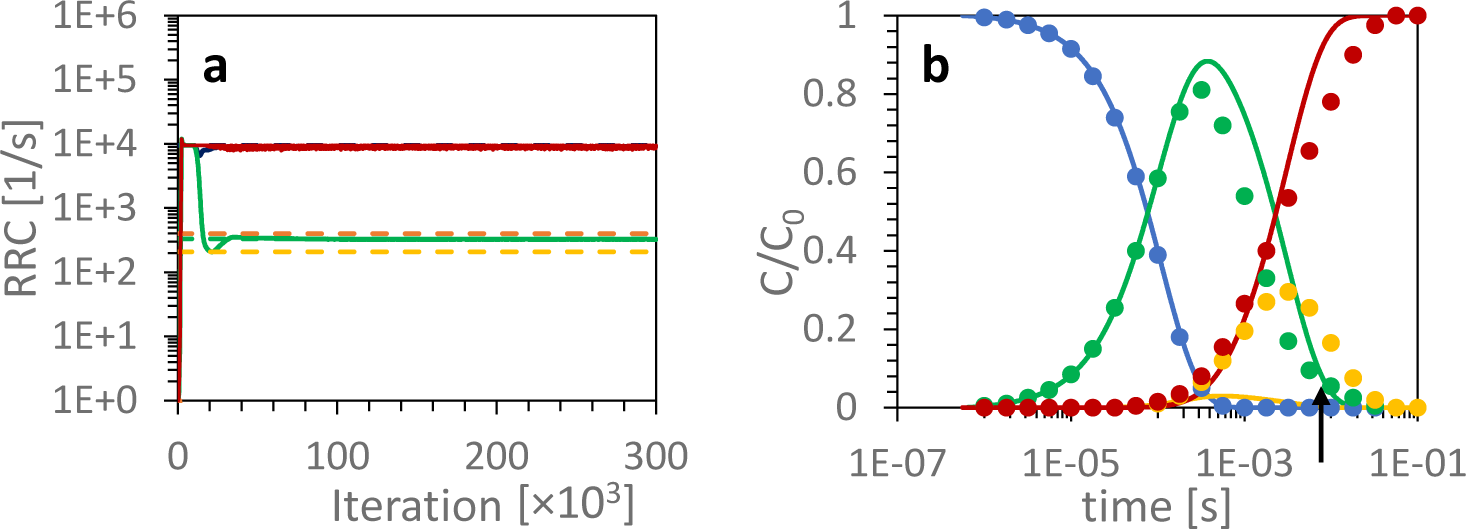
KINNTREX predictions of the RRCs and the concentration profiles of the intermediates retrieved from time-dependent DED maps. (**a**) RRCs vs iteration number for the DE_S_ simulation with lower and upper range limits of 84 and 9500 for 3 RRCs (*k*_*1*_, *k*_*3*_, and *k*_*5*_), respectively. predicted RRCs include *k*_*1*_ (dark blue solid line), *k*_*3*_ (green solid line) and *k*_*5*_ (red solid line). Ground truth RRCs include *k*_*1*_ (dark blue dashed line), *k*_*3*_ (green dashed line), *k*_*-3*_ (yellow dashed line), and *k*_*4*_ (orange dashed line). (**b**) Relative predicted concentration profiles of the intermediates (*C*_*1*_ – blue solid lines, *C*_*2*_ – green solid line, *C*_*3*_-yellow solid line, and *C*_*d*_ – red solid line) for the candidate presented in (**a**) along with ground truth (circles). The colors of the ground truth concentration profiles match the corresponding colors of the predicted concentration profiles. The concentrations are calculated with the RRCs extracted from the last iteration. The upper limits for the RRCs were set to the sum of the relaxation rates, while the lower limit was set to the minimum of the relaxation rates. The arrow in (**b**) indicates the second transient along the concentration profile of I_2_ for the simulated dead-end mechanism.

